# Alcohol Sipping Patterns, Personality, and Psychopathology in Children: Moderating Effects of Dorsal Anterior Cingulate Cortex (dACC)

**DOI:** 10.1101/2024.01.15.575762

**Authors:** Ana Ferariu, Alexei Taylor, Hansoo Chang, Fengqing Zhang

## Abstract

Alcohol, the most consumed drug in the United States, is associated with various psychological disorders and abnormal personality traits. Despite extensive research on adolescent alcohol consumption, the impact of early alcohol sipping patterns on changes in personality and mental health over time remains unclear. There is also limited information on the latent trajectory of early alcohol sipping, beginning as young as 9-10 years old. The dorsal anterior cingulate cortex (dACC) is crucial for cognitive control and response inhibition. However, the role of dACC remains unclear in the relationship between early alcohol sipping and mental health outcomes and personality traits over time. Utilizing the large data from the Adolescent Brain and Cognitive Development study, we aim to comprehensively examine the longitudinal impact of early alcohol sipping patterns on psychopathological measures and personality traits in adolescents, filling crucial gaps in the literature. We identified three latent alcohol-sipping groups, which demonstrate distinct personality traits and depression score trajectories. Bilateral dACC activation during the stop-signal task moderated the effect of early alcohol sipping on personality and depression over time. Additionally, bidirectional effects were observed between alcohol sipping and personality traits. This study provides insights into the impact of early alcohol consumption on adolescent development.

## 1. Introduction

### 1.1 The Effects of alcohol use on brain and behavior

Alcohol is the most consumed drug in the United States while alcohol use disorder (AUD) is one of the most prevalent psychiatric disorders (McHugh & Weiss, 2019). Studies have demonstrated that the intake of alcohol, acutely or chronically, negatively impacts our cognitive functioning due to accelerated, abnormal changes in gray and white matter volume in the frontal and temporal lobes, regions responsible for higher-order cognitive control and learning and memory, respectively (George et al., 2005; Lees et al, 2020; Pfefferbaum et al., 2018). Under the influence of alcohol people are more likely to show impairment in planning and more likely to show increased impulsivity, anger, friendliness, anxiety, and confusion, as well as the sensation of light-headedness (George et al., 2005). AUD patients score lower on visual and verbal working memory tasks and are more likely to develop comorbidity with other substance abuse (Fama et al., 2021). Alcohol has a very strong effect on suppressing the neural networks that are involved in memory formation and storage. When people drink, the prefrontal cortex (PFC) and top-down inhibition are reduced and therefore, habitual, and impulsive behaviors increase for a short period of time. Large amounts of alcohol and constant consumption boost the changes in the neural circuits that underlie impulsive and habitual behaviors, and these changes are projected outside the time when people are drinking. Thus, higher impulsivity and habitual behavior are observed among chronic drinkers, even when they are not under the influence of alcohol (Elvig et al., 2021).

### 1.2 The onset of alcohol use

Early alcohol exposure has an increased risk in developing future alcohol use problems and has a negative impact on the activation of multiple brain regions responsible for reward response, inhibitory control and other cognitive processes (Chaarani et al., 2022). Earlier use of alcohol (9- 10 years old) is associated with increased risk of binge drinking in high school (Jackson et al., 2015; May et al., 2022). Earlier age of drinking onset is associated with alcohol dependence later in life (Grant, 1998) and psychopathology symptoms (Skogen et al., 2016). Children coming from a family with history of AUD are more likely to have reduced anticipatory reward responding (Martz et al., 2022), and reward sensitivity features can influence the degree of alcohol consumption across development (Watts et al., 2023). On the other hand, adolescents who are less likely to sip alcohol present higher self-esteem, behavioral regulation and increased liking of school (May et al., 2022). Multiple studies have examined the relationship between psychopathology or personality traits and alcohol use in adolescents, showing significant results cross-sectionally or longitudinally (Dyer et al., 2019; Fernie et al., 2013; Stautz & Cooper, 2013; Watts et al., 2023). Other studies have explored the latent trajectory of alcohol consumption measured in drinks from mid-adolescence to adulthood, but there is limited information on the latent trajectory of early alcohol sipping, beginning as young as 9-10 years old (Aiken et al., 2022; Chassin et al, 2004; Skogen et al., 2016; van der Vorst et al., 2009; Windle, 2020; Yuen et al., 2020). More research is needed to address this gap, especially by utilizing big data from a large-scale study with rich information collected on alcohol sipping, personality traits, and mental health outcomes over time.

There are lots of factors impacting alcohol initiation and consumption across developmental stages which can be highly influenced by psychopathology and personality traits. Children with anxiety and depressive disorders are more likely to experiment with substance use (Klein et al., 2022). Impulsivity-related traits, such as positive urgency and sensation seeking are positively associated with alcohol consumption, while both negative and positive urgency are associated with problematic alcohol use in older adolescents (Stautz & Cooper, 2013). The associations between behavioral inhibitory system (BIS) traits, behavioral activation system (BAS) traits and alcohol sipping are significant in adolescents and consistent across stages of youth development (Watts et al., 2023). In particular, BAS reward responsiveness is a significant factor for high-risk behaviors, such as the use of drugs (Assari et al., 2020), while weak inhibitory control is associated with alcohol misuse in adolescents (Lopez-Caneda et al., 2013). The impact of early alcohol sipping on psychopathological measures and personality traits over time needs to be further evaluated. Inconsistent findings exist regarding the reciprocal effects of alcohol consumption and impulsivity traits in adolescents, with varying cohorts and ages demonstrating bidirectionality or unidirectionality (Farley & Kim-Spoon, 2015; Kaiser et al., 2015; Kaiser et al.,2016; Malmberg et al., 2013; Nos, 2014). More work is needed to investigate the potential bidirectional relationship between alcohol sipping and psychopathology or personality traits, evolving from childhood to early adolescence.

### 1.3 The role of the dACC in inhibition

The dorsal anterior cingulate gyrus (dACC) plays a very important role in cognitive control and response inhibition, as well as in reward-based learning, decision-making, motor functions, emotion and autonomic functions (Bush et al., 2000; Picard & Strick, 2001; Smith et al., 2009; Somerville et al., 2006). In adolescence, there is a notable restructuring of ACC connectivity (Fair et al., 2007; Ho et al., 2017; Kelly et al., 2009; Power et al., 2010).

Specifically, during childhood, dACC connectivity tilts towards the central executive network (CEN), but as individuals transition into adulthood, this connectivity shifts towards the salience network (SN) (Fair et al., 2009; Ho et al., 2017, Manza et al., 2016; Power et al., 2010; Sole-Padulles et al., 2016; Uddin et al., 2011).

The stop-signal task (SST) is one of the most widely used tasks to analyze response inhibition and inhibitory control (Lipszyc & Schachar, 2010). The SST also presents behavioral measures regarding motor anticipation under cognitive control (Boen et al., 2022). Lesion studies involving dACC showed compromised response inhibition abilities, especially in the context of adapting to a reduced reward amount (Heilbronner & Hayden, 2016; Rushworth et al., 2003; Shima & Tanji, 1998; Williams et al., 2004). Dysfunction in the dACC mirrors cognitive disruptions, particularly in the areas of impaired response selection and processing information that demands cognitive effort (Yücel et al., 2003).

Moreover, a significant body of evidence linked dACC dysfunction to various psychological disorders (Benes, 1993; Critchley et al., 2003; Ridderinkhof et al., 2004; Yücel et al., 2003) as well as to normal and abnormal personality traits (Canli et al., 2004; Gray et al., 2005; Kumari et al., 2004; Whittle et al., 2006; Pujol et al., 2002). Variability in dACC activation in response to emotional stimuli can be influenced by personality traits or mood states (Canli et al., 2004). Poor inhibitory control is a characteristic of OCD (Woolley et al., 2008) and AUD (Goudriaan, 2006) and extended stop-signal reaction times (SSRTs) in high-risk adolescents can predict substance use disorders, including AUD (Nigg et al., 2006). The link between AUD and dACC function becomes most apparent during cognitively demanding tasks (Murray et al., 2022). Nevertheless, the precise influence of dACC activity on psychopathology and personality traits over time remains somewhat ambiguous in adolescent development. It is uncertain whether dACC activity during childhood positively or negatively shapes the development of psychopathology or personality traits, given its evolving connectivity throughout adolescence and into adulthood. Additionally, the role of dACC remains unknown in the relationship with early alcohol sipping behavior and mental health outcomes and personality traits over time and therefore needs to be further examined.

### 1.4 Present study

To address the questions outlined above, our study commences with an examination of the latent trajectory of early alcohol sipping using data from the Adolescent Brain and Cognitive Development (ABCD) cohort. This initial exploration provides valuable insights into representative patterns of alcohol sipping behavior as individuals transition from childhood to adolescence. Leveraging latent class mixed models, which can analyze unobserved longitudinal patterns and handle missing data, enhances the robustness of our analysis. Subsequently, considering the longitudinal and nested structure of the ABCD data, we evaluate the impact of these latent alcohol sipping classes on mental health outcomes and personality traits over time. Additionally, we investigate the potential moderating role of the dorsal anterior cingulate cortex (dACC), crucial in response inhibition and during development, in shaping the effects of the latent trajectory of alcohol sipping on psychopathology and personality traits. In our exploratory analyses, we aspire to investigate the bidirectional relationships among alcohol use, personality traits, and mental health outcomes. In essence, our study aims to comprehensively examine the longitudinal impact of early alcohol sipping patterns on psychopathological measures and personality traits in adolescents, addressing a crucial gap in the literature. This endeavor is particularly notable for its utilization of extensive data from a population-based study, offering rich information on alcohol sipping, personality traits, and mental health outcomes over time.

## 2. Methods

### 2.1 Participants

We used data from the 5.0 release of the Adolescent Brain Cognitive Development (ABCD) study, the largest longitudinal study of brain development and youth health in the United States. Data was available at three time points (baseline, 2-year and 4-year) for the personality traits outcomes and at five time points (baseline, 1-year, 2-year, 3-year and 4-year) for the mental health outcomes and the alcohol sipping variable. The baseline ABCD cohort was represented by 11,868 participants (52% white, 52% males) coming from 9807 unique families across 22 sites with mean age of 119 months (9-10 years old, SD = 7.5).

### 2.2 Measures

#### 2.2.1 Alcohol exposure

To assess early alcohol exposure, we used data measured by the iSay Sip Inventory (Jackson et al., 2015), including questions regarding age of alcohol use onset, number of sips and type of alcohol ingested. The variable of interest in our analysis was the number of alcohol sips reported by the participants. The baseline value for the number of sips included the number of all sips prior to the study, while the sipping variable at the yearly follow-ups included the number of sips in the past year.

#### 2.2.2 Personality traits

Impulsivity traits and behavioral inhibitory/approach systems (BIS/BAS) scores were measured using the UPPS-P for Children Short Form (Watts et al., 2020) and BIS/BAS scales (Carver & White, 1994; Pagliaccio & Luking, 2016), respectively. We used impulsivity sub- scores for negative urgency (NU), positive urgency (PU), lack of planning (LPlan), lack or perseverance (LPer), sensation seeking (SS) and BIS/BAS sub-scores for BIS, BAS fun, BAS drive and BAS reward responsiveness. Impulsivity sum scores and BIS/BAS sum scores represented independent traits. Impulsivity sum scores were set on a scale ranging from 4-16, while BIS/BAS sum scores were measured on a scale ranging from 0-12, with lower scores implying lower propensity for the specific trait. The UPPS-P and BIS/BAS scales assess impulse action and response inhibition (Bari & Robbins, 2013; Green et al., 2023)

#### 2.2.3 Mental health outcomes

Mental health outcomes were measured by the parent-reported Child Behavior Checklist (CBCL) and included sum scores for depression, anxiety, obsessive-compulsive disorder (OCD), stress and comorbid anxiety and depression, based on DSM-5 diagnosis and criteria (Achenbach et al., 2003, 2007; Barch et al., 2021). CBCL sum scores ranged from 50 to 100.

Mania sum scores were measured by the Parent General Behavioral Inventory (PGBI) - Mania and ranged from 0 to 30 (Barch et al., 2021; Youngstrom et al., 2008).

#### 2.2.4 dACC Activation during the SST

We used task functional magnetic resonance imaging (task-fMRI) data from the SST to measure the dACC activation at the baseline visit for the two contrasts responsible for inhibitory control (correct-stop-vs-incorrect-stop and correct-stop-vs-correct-go) (Charaani et al., 2021).

The correct-stop-vs-incorrect-stop SST captured the dACC neural activity in the contrast between successful inhibitory control and unsuccessful/failed inhibitory control, while the correct-stop-vs-correct-go SST contrast captured the dACC neural activity between successful inhibitory control and successful execution of a planned response. The neuroimaging variable was measured as the mean beta-weight for all contrasts in cortical Destrieux ROI right and left hemisphere separately and for each contrast. The neuroimaging data was pre-processed by the ABCD study using FreeSurfer software (Fischl, 2012), while the pipeline can be found elsewhere (Hagler et al., 2019), and we performed additional quality control based on the critiques and recommendations of Bissett et al. (2021) and Garavan et al. (2022).

### 2.3 Statistical Analysis

Our analysis was divided into five parts: (i) preliminary analysis for exploring alcohol sipping patterns over time; (ii) identifying potential latent trajectory of alcohol sipping over time; (iii) the effects of the identified latent alcohol sipping classes on changes in personality traits and mental health outcomes over time (iv) the role of dACC as a moderator in the relationship between latent alcohol sipping patterns and personality traits or mental health outcomes; and (v) exploring the bidirectionality between alcohol sipping and personality traits and mental health scores over time. All statistical analyses were performed using R 4.3.0 (R Core Team, 2023).

#### 2.3.1 Preliminary Analysis for Alcohol Sipping Behavior

We classified the number of sips into three categories at each time point: no-sip (zero sips), low-sip (1 or 2 sips) and high-sip (at least 3 sips), similarly to the categorization of Watts et al. (2023) . The outliers of the number of alcohol sips were truncated to the mean plus three standard deviations. We calculated the percentage of participants in each category and at each time point (baseline, 1-year follow-up, 2- year follow-up, 3- year follow-up, 4- year follow-up).

### 2.3.2 Latent Trajectories of Alcohol Sipping in Adolescents

We conducted latent class mixed models using the lcmm package in R (Proust-Lima et al., 2023) with the goal of detecting latent classes representative of the alcohol sipping behavior over time. We used data from 1-year follow-up to 4-year follow-up, allowing us to capture the sipping behavior from baseline up to the current moment. Alcohol sipping measured at baseline was not included in the analysis because it represented the number of all sips prior to the study. We ran a total of four models using the beta link function and fitting one, two, three and four classes with the intention of comparing the four models and choosing the one that would represent our data the best, based on the lowest Akaike Information Criterion (AIC) and Bayesian Information Criterion (BIC) (Akaike, 1973; Schwarz, 1978).

### 2.3.3 The Effects of Latent Alcohol Sipping Patterns on Personality Traits and Mental Health Outcomes

After identifying the latent classes related to the number of alcohol sips over time, we examined how these latent classes differentially impact the trajectory of personality traits and mental health outcomes over time. Multilevel modeling was conducted for each of the outcomes separately using the lme4 package in R (Bates et al., 2015), with time as a predictor and a random effect structure involving subject ID, family ID and site ID. Given that personality data was collected only at three time points, a linear term for time was utilized in the analysis. For mental health outcomes collected over five time points, both linear and quadratic time effects were examined. When needed, the outcomes were rescaled and transformed to meet the normality assumption. Restricted maximum likelihood was used to estimate model parameters and to test the significance of random effects. Model selection criteria such as AIC and BIC were used to select the best-fitting model. For mania and CBCL scores, we conducted compound Poisson generalized linear mixed models using the cplm package in R (Zhang, 2013) to account for their zero-inflated distribution. After identifying the optimal model for each outcome, we added an interaction term between the function of time and the latent classes to determine whether the trajectory of the outcome over time differed significantly by the latent alcohol sipping groups. We controlled for covariates including age at baseline (measured in months), sex assigned at birth by parents (1 male, 0 otherwise), race assigned at birth by parent (1 white, 0 otherwise), family history of alcoholism (1 if any family member reported AUD, 0 otherwise) and socio-economic status (SES) measured as the Area Deprivation Index (ADI) on a scale from 1 to 125.

#### 2.3.4 The Role of dACC in the Relationship with the Latent Trajectories on Personality Traits and Mental Health Outcomes

Additionally, for each of the outcomes found to show significantly different trajectories for different latent alcohol sipping groups, we tested whether the left and right dACC activations during the SST at baseline further moderated the impact of the latent alcohol sipping groups on the outcomes. We specifically examined two SST contrasts, correct-stop-vs-correct-go and correct-stop-vs-incorrect-stop, to account for response inhibition.

#### 2.3.5 Exploring Bidirectional Associations between Alcohol Sipping and Personality Traits and Mental Health Outcomes

We first explored the bidirectionality between alcohol sipping and personality traits and mental health outcomes using generalized linear mixed models at each of the three time points (baseline, 2-year follow-up, 4-year follow-up) separately. We examined whether alcohol sipping measured at one time point predicted each of the outcomes measured at the following time point and vice versa. All models included a random effect structure involving family ID, site ID and covariates related to demographics such as sex, race, age at baseline, SES and family history of alcoholism, and baseline scores. For the outcomes found to have significant bidirectional association with alcohol sipping, we further conducted a random intercept cross-lagged panel model (RI-CLPM) using the lavaan package in R (Rosseel, 2012) to test the existence of significant simultaneous bidirectionality by jointly modeling their trajectories over time. A random intercept was needed to account for stable, trait-like differences between individuals in the initial levels of alcohol sipping and sum scores and for unobserved heterogeneity (Mulder & Hamaker, 2021; Mund et al., 2021). We included time-invariant person level characteristics (i.e. socio-economic status, age at baseline, family history of alcoholism, sex and race) at the random intercept level (Mulder & Hamaker, 2021). We decomposed the observed structure into 3 components: grand means, between-person components and within-person components. Since the data was non-normally distributed, we used the “MLR” estimator in the lavaan function to obtain robust standard errors (Yuan and Bentler, 2000), as well as a full information maximum likelihood (FIML) for missing data (Lee & Shi, 2021). Model goodness-of-fit was achieved if the Comparative Fit Index (CFI) and the Tucker-Lewis Index (TLI) were greater than 0.90 and the Root Mean Squared Error of Approximation (RMSEA) was lower than 0.06 (Bentler, 1990).

## 3. Results

### 3.1 Descriptive Statistics of Alcohol Sipping Behavior Over Time

At baseline 77.5% of the participants have never had an alcohol sip in their lifetime, while 16.2% had either 1 or 2 sips in their lifetime and 6.3% had at least 3 alcohol sips in their lifetime (Fig. 1). At the 1-year follow-up, 6.8% and 2.8% of the participants recorded having 1-2 alcohol sips or more than 3 sips in the past year, respectively. 90.4% of the participants have still reported no alcohol sipping. However, starting from Year 1, the percentage in the “no-sip” category decreased over time with 83.0% at the 4-year follow-up, while the percentage of the participants in the “high-sip” category increased consistently over time with 7.9% at the 4-year follow-up. The percentage of participants in the “low-sip” category remained stable around 6.7% (+/- 0.1%) until the 3-year follow-up. However, this category showed a 1.3-fold increase at the 4-year follow-up reaching 9%. In general, the adolescents exhibited a rising trend in the quantity of alcohol sips consumed over the course of four years.

**Fig. 1.**
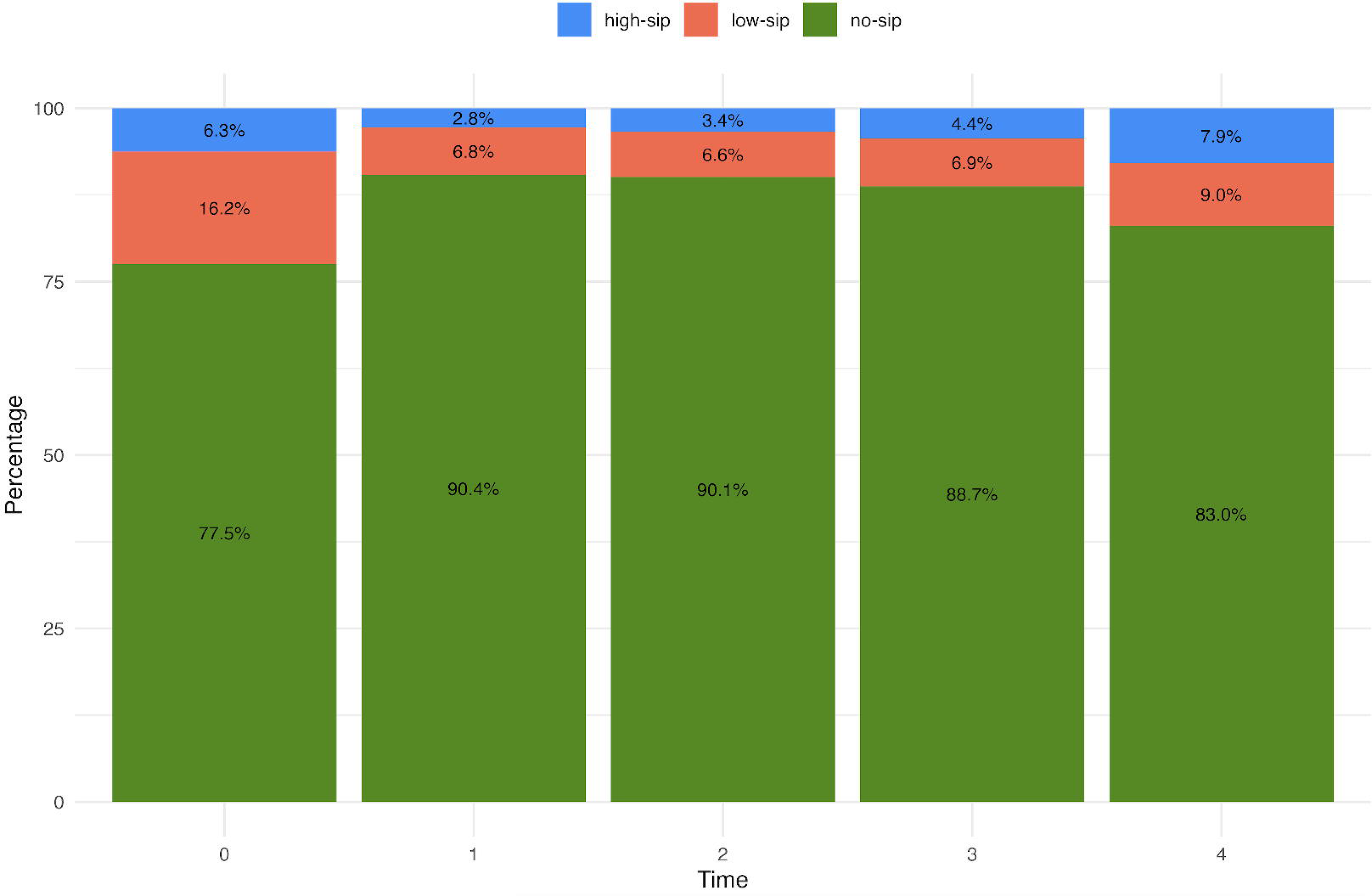
The percentage of participants in each sipping category at each time point

### 3.2 Latent Trajectories of Alcohol Sipping in Adolescents

Our analysis results (**Table S1**) show that the model with 3 latent classes fitted the data best with the lowest AIC and BIC (Akaike, 1973; Schwarz, 1978). The identified three latent alcohol sipping groups are shown in **Fig. 2**. Most of the participants belonged to latent class 2 (84.22%, n = 9700), whose trajectory over time was almost constant around zero sips (no-sip group). Latent class 1 (low-sip group), represented by the smallest percentage of participants (5.34%, n = 615) showed a decreased trajectory of alcohol sips over time, even though this group of participants represented the highest mean of number of sips between baseline and 1-year follow-up. The average curve for latent class 3 (high-sip group, 10.44% participants, n = 1202) started below the average number of sips of latent class 1 but presented a steady increase in the mean number of sips over time.

**Fig. 2.**
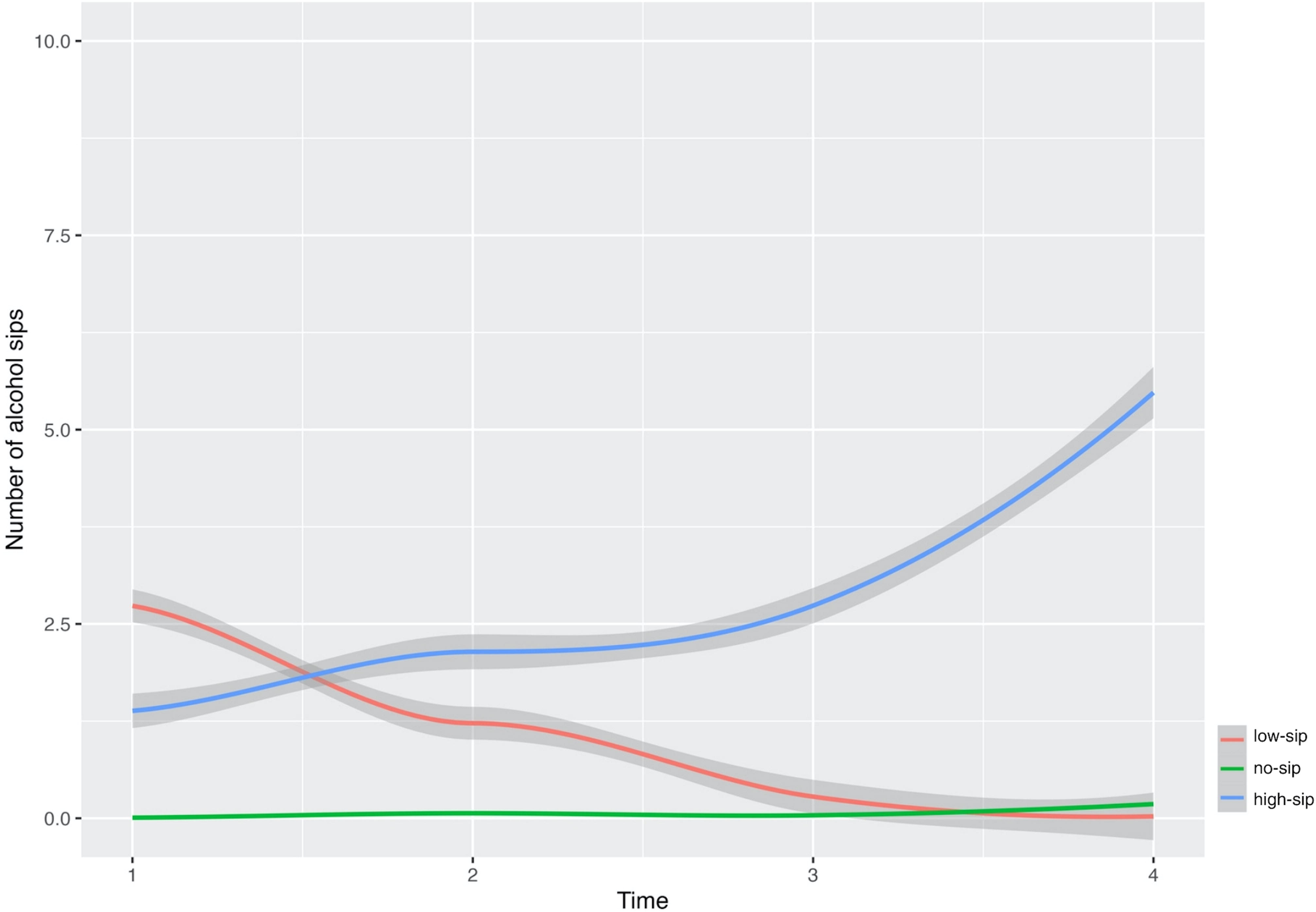
**The average curve for the latent classes representing the sipping behavior over time in the ABCD cohort and the corresponding standard error.**

### 3.3 The Effects of Latent Alcohol Sipping Trajectories on Personality Traits and Mental Health Outcomes

As shown in **Table 1**, the two-way interaction terms between the latent alcohol sipping classes and time were statistically significant for several personality trait outcomes. Compared to the high-sip group, the no-sip group had on average a slower rate of increase in negative urgency (b= -0.11, p < 0.001), positive urgency (b = -0.10, p < 0.001), lack of perseverance (b = - 0.05, p < 0.001), sensation seeking (b = - 0.16, p < 0.05), BIS scores (b = -0.10, p < 0.001), BAS reward responsiveness scores (b = -0.17 p < 0.01), BAS fun scores (b = -0.11, p < 0.001) and BAS drive scores (b = -0.03, p < 0.05). In other words, the effects of time on these personality traits were weaker for the group with consistently low or no sipping compared to the group showing an increase in the number of sips over the four-year period.

**Table 1.** Summary table for the personality trait outcomes.

|  | NU | PU | LPlan | LPer | SS | BIS | BAS RR | BAS fun | BAS drive |
| --- | --- | --- | --- | --- | --- | --- | --- | --- | --- |
| (Intercept) | **1.80\*** | **1.21\*** | **0.75\*** | **0.97\*** | **7.92\*** | **2.47\*** | **8.51\*** | **2.13\*** | **1.14\*** |
| Time | **0.05\*** | **0.04\*** | 0.10 | **0.07\*** | **0.15\*** | 0.03 | **-0.51\*** | **-0.07\*** | 0.02 |
| lc1 | 0.05 | 0.02 | 0.15 | 0.02 | -0.06 | 0.05 | 0.18 | 0.01 | -0.00 |
| lc2 | **-0.08\*** | **-0.10\*** | **-0.33\*** | -0.03 | **-0.78\*** | 0.02 | -0.01 | **-0.07\*** | -0.03 |
| Family History of Alcoholism | **0.04\*** | 0.02 | **0.15\*** | **0.06\*** | 0.07 | **0.02\*** | -0.02 | 0.01 | **-0.03\*** |
| Sex | **0.09\*** | **0.15\*** | **0.33\*** | **0.06\*** | **0.62\*** | **-0.20\*** | **0.14\*** | **0.06\*** | **0.09\*** |
| Race | **0.06\*** | **0.16\*** | **-0.15\*** | **0.04\*** | **-0.22\*** | **-0.04\*** | **0.22\*** | **0.04\*** | **0.09\*** |
| SES | 0.02 | **0.10\*** | -0.030 | **0.05\*** | **-0.23\*** | **-0.05\*** | 0.01 | 0.03 | **0.04\*** |
| Age | **0.00\*** | -0.00 | 0.002 | 0.00 | **0.02\*** | **0.00\*** | 0.00 | -0.00 | **0.00\*** |
| time:lc1 | **-0.09\*** | **-0.11\*** | **-0.212\*** | **-0.06\*** | **-0.30\*** | -0.02 | -0.12 | **-0.10\*** | -0.03 |
| time:lc2 | **-0.11\*** | **-0.10\*** | 0.009 | **-0.05\*** | **-0.16\*** | **-0.10\*** | **-0.17\*** | **-0.11\*** | **-0.03\*** |
| Notes: * $p < 0.05$. lc1 and lc2 are dummy variables, representing high-sip (lc1=0, lc2=0), low-sip(lc1=1, lc2=0) and no-sip (lc1=0, lc2=1). | | | | | | | | | |

Additionally, the low-sip group had on average a slower rate of increase over the 4-year follow-up in negative urgency (b = - 0.09 p < 0.01), positive urgency (b = - 0.11, p < 0.01), lack of planning (b = -0.21 p < 0.05), lack of perseverance (b = -0.06 p < 0.05), sensation seeking (b = - 0.30, p < 0.01 and BAS fun scores (b = -0.10 p < 0.001) compared to the high-sip group. Findings on the mental health outcomes are shown in **Table 2**. Depression score was the only outcome showing significantly different trajectories over time for different latent alcohol sipping groups. The significant two-way interactions between quadratic time term and latent alcohol sipping classes (b = -0.039, p < 0.05 for time^2^*lc1 and b = -0.022, p < 0.05 for time^2^*lc2) suggest that depression scores for the no-sip and low-sip groups followed an inverted U-shape curve, slowly increasing during the first two years and decreasing afterwards while the high-sip group had a monotonically increasing pattern of depression scores over the four years. The no- sip group on average had the lowest depression scores over time. At the 4-year follow-up, the high-sip group showed the highest depression scores, followed by the low-sip group and then the no-sip group.

**Table 2.** Summary table for mental health outcomes.

|  | Anx | Dep | AnxDep | OCD | Stress | Mania |
| --- | --- | --- | --- | --- | --- | --- |
| (Intercept) | **-1.62\*** | -0.81 | **-1.03\*** | **-0.93\*** | **-2.27\*** | **-4.19\*** |
| Time | **0.2\*** | 0.16 | **0.09\*** | **0.31\*** | 0.06 | **0.21\*** |
| lc1 | 0.11 | 0.06 | 0.15 | **0.25\*** | 0.19 | 0.19 |
| lc2 | **0.17\*** | -0.03 | 0.10 | 0.11 | 0.02 | -0.03 |
| I(time^2) | **-0.06\*** | -0.02 | **-0.03\*** | -0.10 | -0.00 | **-0.11\*** |
| Family History of Alcoholism | **0.68\*** | 0.58 | **0.76\*** | 0.70 | **0.75\*** | **0.37\*** |
| Sex | **0.35\*** | 0.10 | 0.01 | 0.04 | **0.25\*** | **0.09\*** |
| Race | -0.01 | 0.03 | **-0.13\*** | **-0.23\*** | **0.11\*** | **0.54\*** |
| SES | **0.15\*** | 0.16 | **0.11\*** | 0.06 | **0.32\*** | **0.56\*** |
| Age | **0.01\*** | 0.00 | 0.00 | **0.01\*** | 0.00 | 0.00 |
| time:lc1 | 0.00 | 0.08 | 0.04 | -0.01 | -0.01 | 0.05 |
| time:lc2 | -0.08 | -0.00 | -0.02 | -0.00 | -0.07 | -0.04 |
| lc1:I(time^2) | -0.01 | **-0.04\*** | -0.01 | 0.00 | -0.02 | -0.03 |
| lc2:I(time^2) | 0.00 | **-0.02\*** | -0.02 | -0.02 | -0.01 | -0.01 |
| Notes: * $p < 0.05$. lc1 and lc2 are dummy variables, representing high-sip (lc1=0, lc2=0), low-sip (lc1=1, lc2=0) and no-sip (lc1=0, lc2=1). | | | | | | |

### 3.4 dACC Moderates the Effect of Latent Alcohol Sipping Groups on Personality Traits and Depression Scores

#### Correct stop versus incorrect stop contrast

When examining the contrast of dACC activation between correct stop and incorrect stop measured at baseline, we found significant 3-way interaction effects between time, the no-sip vs. high-sip groups and left dACC for lack of perseverance (b = 0.10, p < 0.05), sensation seeking (b = 0.44, p < 0.05) and BAS fun (b = 0.17, p < 0.01). Moreover, sensation seeking also showed significant 3-way interaction between time, the no-sip vs. high-sip groups and the right dACC (b = -0.49, p < 0.05), as well as BAS fun (b = -0.15, p < 0.01) and BAS drive (b = -0.12, p < 0.05). The full output for the moderation analyses can be found in the **Tables S2-S11**. **Fig. 3a and 3b** depict the trajectory of sensation seeking scores over time for the left and right dACC separately. To illustrate the moderating effect of dACC on the differences in sensation seeking between the no-sip and high-sip groups over time, we graphed the patterns for low, average and high levels of dACC activation separately in **Fig. 3**. The low and high levels of dACC activation were computed as the mean dACC activation minus or plus one standard deviation, respectively. As shown in **Fig. 3a**, the differences in sensation seeking between the no-sip and high-sip groups over time were larger for participants with lower left dACC activation, suggesting that the left dACC activation significantly moderated the effect of the no-sip vs. high-sip groups on sensation seeking. Similarly, we observe the moderating effect of right dACC, as indicated by a larger difference in sensation seeking between the no-sip and high-sip groups for participants with higher right dACC activation (**Fig. 3b**). Additionally, non-sippers on average consistently reported lower sensation seeking scores across all time points compared to high sippers. Within the no-sip group, individuals with low baseline activation of the left dACC exhibited a decline in sensation seeking scores over time compared to those with high baseline left dACC activation.

**Fig. 3.**
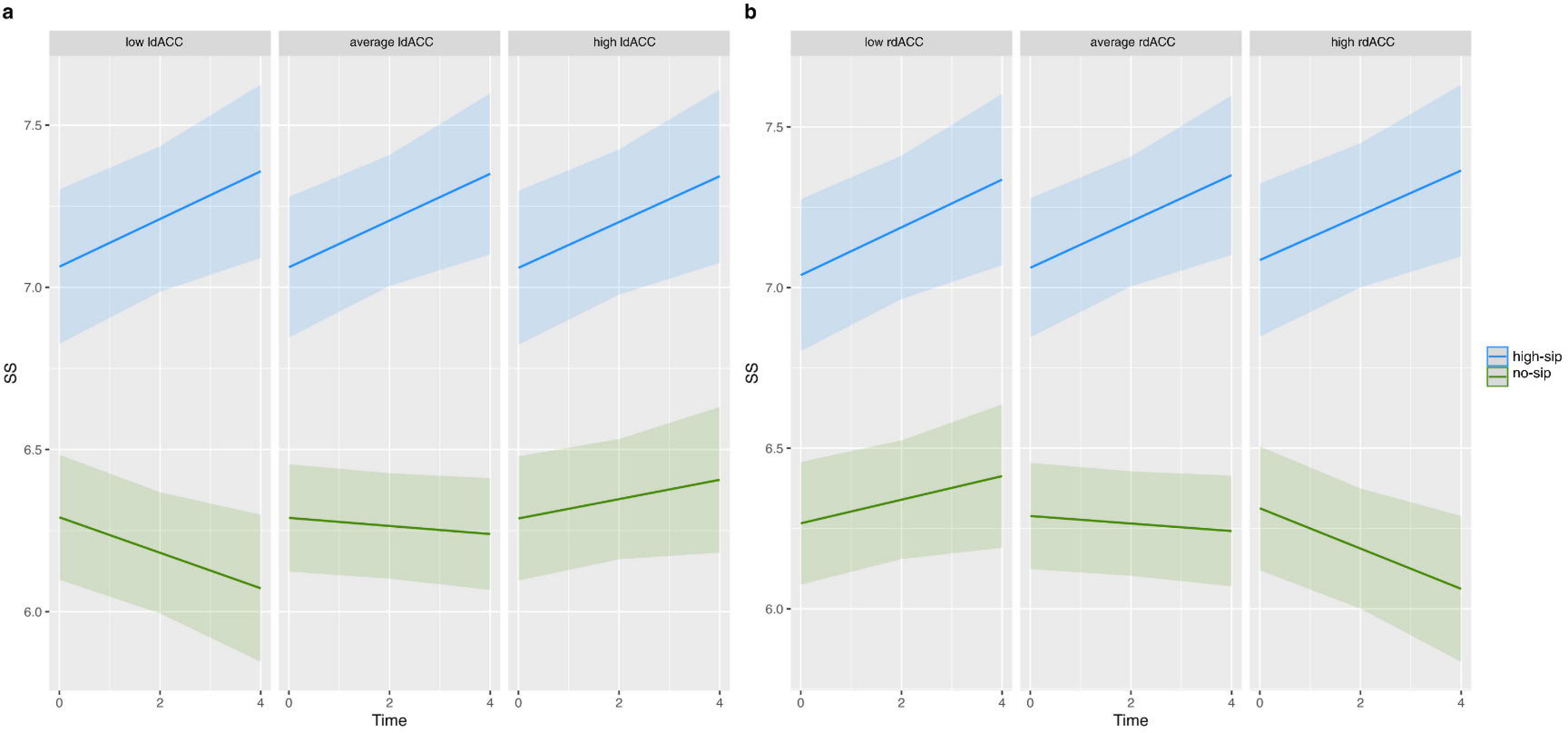
The trajectory of sensation seeking scores for the high-sip vs. no-sip groups for low, average and high neural activity in the (a) left dACC and (b) right dACC in the correct- stop-vs-incorrect-stop SST contrast measured at baseline.

Conversely, when considering baseline right dACC activation, non-sippers with high activation demonstrated a decrease in sensation seeking scores over time, whereas those with low right dACC activation showed a slight increase in sensation seeking scores over time. A similar pattern was present in the trajectory of lack of perseverance scores over time, but the moderating effect of dACC was significant only in the left hemisphere (**Fig. S1**).

The trajectories of BAS fun scores over time for baseline left and right dACC activation during the correct-stop-vs-incorrect-stop SST contrast are shown in **Fig. 4a and 4b**. In alignment with the sensation-seeking results, non-sippers with low left dACC and high right dACC activation at baseline exhibited a more pronounced decline in BAS fun scores over time compared to participants in the high-sip group. This illustrates the effect of latent alcohol sipping groups (no-sip vs. high-sip) on BAS fun trajectory over time was moderated by baseline dACC activation. Moreover, the significant moderating effect of right dACC on the relationship between the latent alcohol sipping groups and BAS drive scores is shown **Fig. S2**. We observe a significantly larger difference in BAS drive scores between the no-sip and high-sip groups over time for participants with higher right dACC activation. The trajectory of BAS drive scores for the non-sippers and high sippers with low right dACC activation were not significantly different. However, the trajectories of BAS drive diverged between the no-sip and high-sip groups over time for participants with higher right dACC activation.

**Fig. 4.**
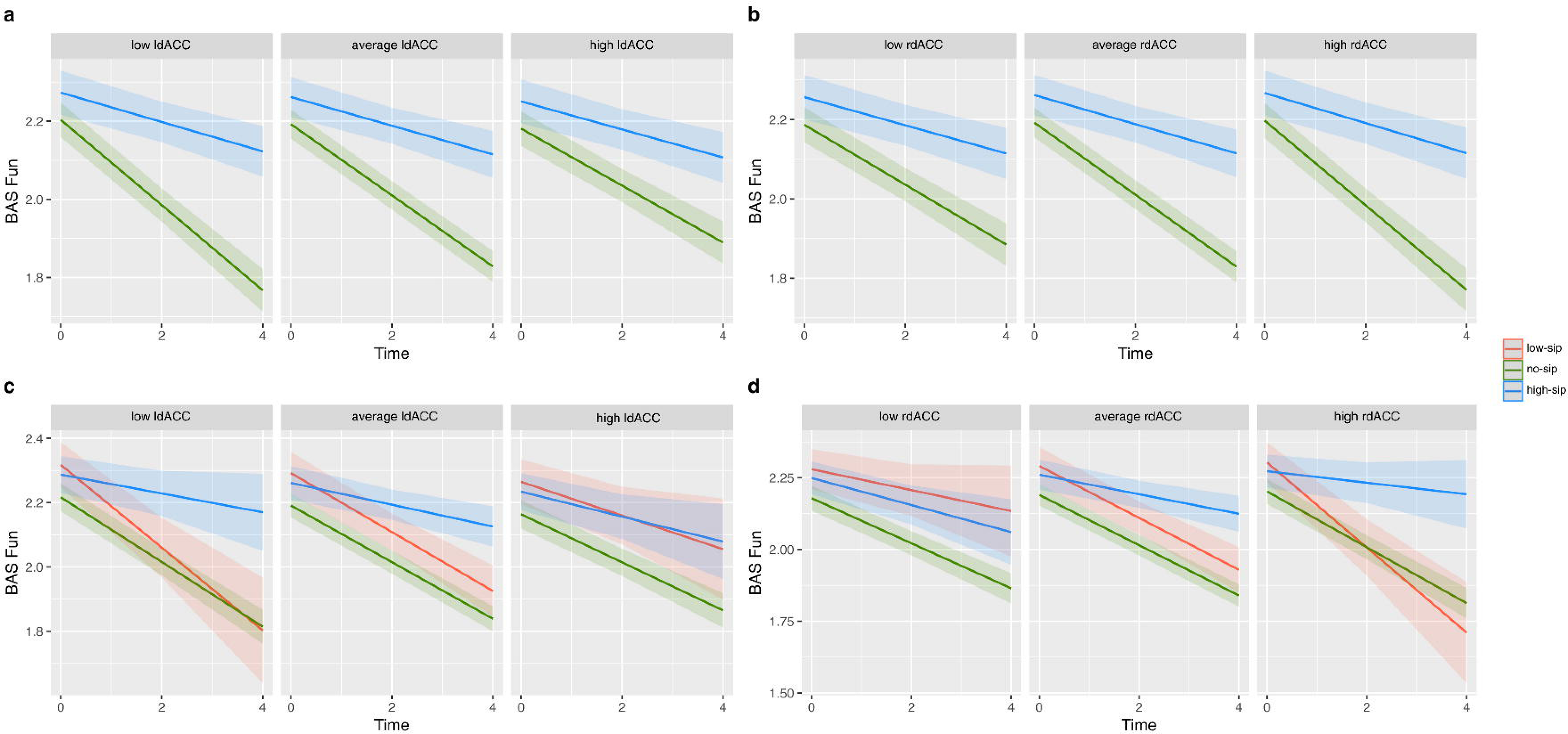
The trajectory of BAS fun scores for each alcohol sipping group for low, average and high dACC neural activity measured at baseline during the correct-stop-vs-incorrect- stop SST contrast in the (a) left hemisphere and (b) right hemisphere, as well as during the **correct-stop-vs-correct-go SST contrast in the (c) left hemisphere and (d) right hemisphere.**

For depression scores, there were significant 3-way interactions between the quadratic time term, the no-sip vs. high-sip groups, and the left dACC (b = -0.13, p < 0.001), and significant 3-way interaction between time, the no-sip vs. high-sip groups, and left dACC (b = 0.35, p < 0.05). This implies the diverging trajectories of depression scores over time between the no-sip and high-sip groups were significantly more separate for participants with higher left dACC activation (**Fig. 5a**). The low, average and high levels of dACC activation were computed based on the tertiles of the dACC activation. The right dACC showed significant interaction effects between the quadratic time term and the no-sip vs. high-sip groups on depression levels (b = 0.08, p < 0.05), suggesting the differences in depression scores over time between the no-sip and high-sip groups were more pronounced for participants with lower right dACC activation (**Fig. 5b**).

**Fig. 5.**
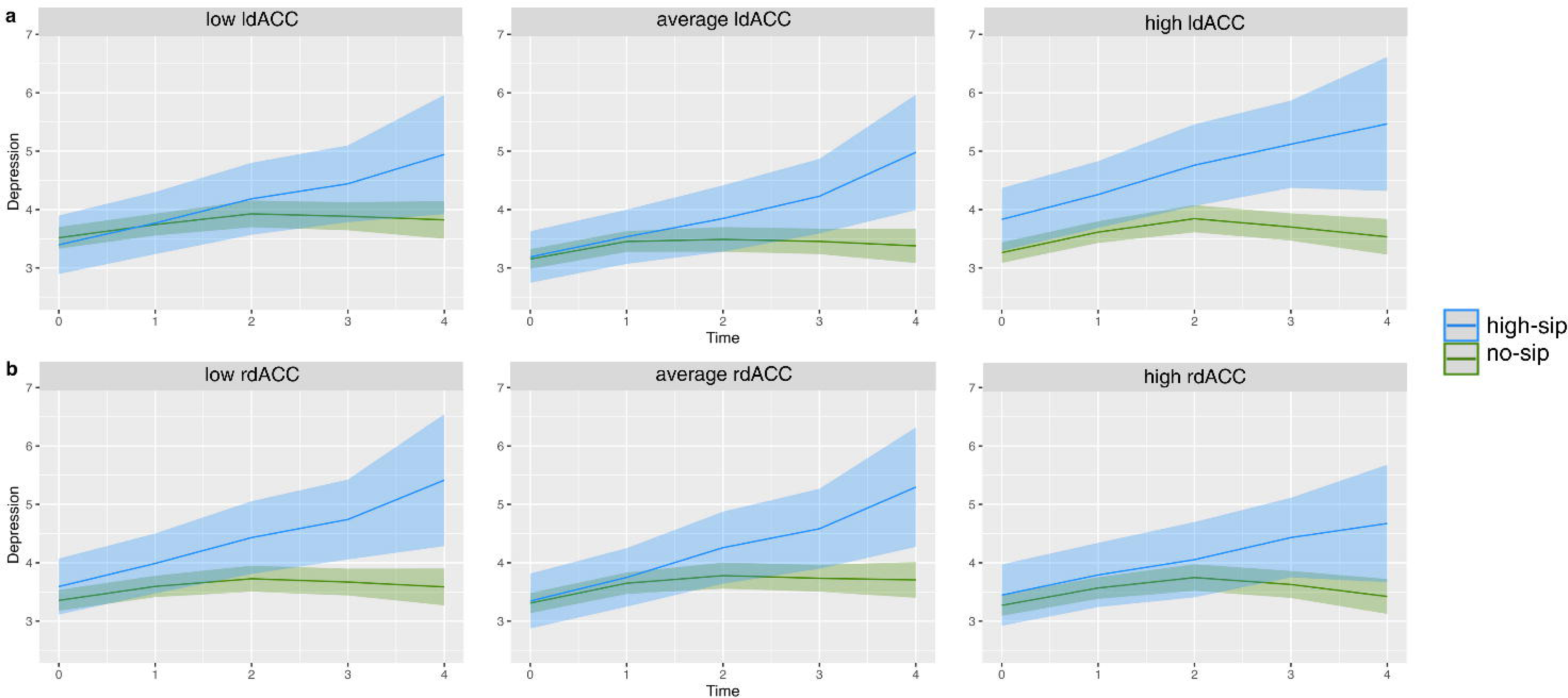
The trajectory of depression scores for the high-sip vs. no-sip groups for low, average and high neural activity in the (a) left dACC and (b) right dACC in the correct- stop-vs-incorrect-stop SST contrast measured at baseline.

#### Correct stop versus correct go contrast

BAS fun scores showed significant 3-way interaction effects between time, the low-sip vs. high-sip groups and left dACC activation (b = 0.47, p < 0.05), as well as for right dACC activation (b = -0.73, p < 0.01). Similarly, there were significant 3-way interactions between time, the no-sip vs. high-sip groups and left dACC (b = 0.16, p < 0.05) as well as for right dACC (b = - 0.13, p = 0.061), but the latter was only borderline significant. The moderating effect of baseline left and right dACC activation during the correct-stop-vs-correct-go SST contrast on BAS fun trajectories over time are illustrated in **Fig. 4c and 4d**. The differences in BAS fun scores between the no-sip and high-sip groups and the differences between the low-sip and high-sip groups over time were larger for participants with lower left dACC activation, suggesting that the left dACC activation significantly moderated the effect of the no-sip and low-sip vs. high sip groups on BAS fun scores (**Fig 4a**). Similarly, we observe the moderating effect of right dACC, as illustrated by a larger difference in BAS fun scores between the low-sip and high-sip groups and between the no-sip and high-sip groups for participants with higher right dACC activation (**Fig. 4d**). Over time, the average of BAS fun scores decreased for all sipping groups.

Bilateral dACC activation was found to significantly moderate the effect of the no-sip vs. high-sip groups on the trajectory of BAS drive over time (b = 0.12, p < 0.05 for left dACC; b = - 0.15, p < 0.01 for right dACC). **Fig. 6a** captures the trajectory of BAS drive scores over time between the no-sip vs. high-sip groups for the left dACC activation. The difference between the BAS drive scores was the largest between the no-sip and high-sip groups when participants showed lower left dACC activation at baseline. Similarly, participants in the no-sip vs high-sip groups with higher right dACC activation also showed a more pronounced difference in BAS drive scores (**Fig 6b**). This implies that the left and right dACC significantly moderated the effect of the no-sip vs. high-sip groups on BAS drive.

**Fig. 6.**
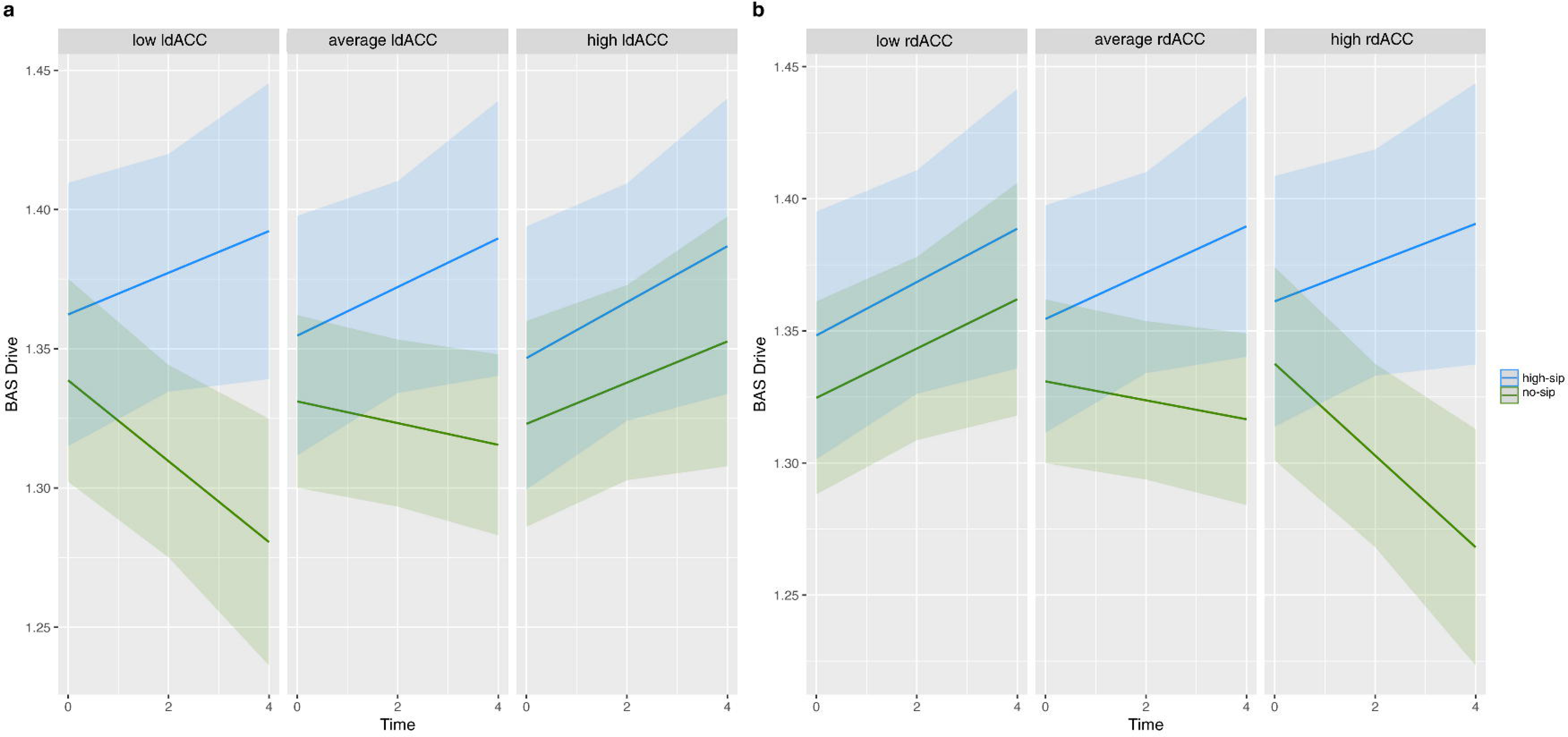
The trajectory of BAS drive scores for the high-sip vs no-sip group for low, average and high neural activity in the (a) left dACC and (b) right dACC in the correct-stop-vs- correct-go SST contrast measured at baseline.

For depression scores, we found significant 3-way interactions between the quadratic time term, the no-sip vs. high-sip groups, and the left dACC (b = - 0.09, p < 0.05), between time, the low-sip vs. high-sip groups, and left dACC (b = -1.92, p < 0.001) and between time, the low- sip vs. high-sip groups and right dACC (b = 2.43, p < 0.001). These results suggest that bilateral dACC activation significantly moderated the effects of the no-sip and low-sip groups on the trajectory of depression levels over time, relative to the high sip group. Additionally, the directionality of the moderation effects on depression levels in the correct-stop-vs-correct-go contrast was consistent with the directionality in the correct-stop-vs-incorrect-stop contrast.

### 3.5 Bidirectional Associations between Alcohol Sipping and Personality Traits and Mental Health Outcomes

There was evidence of significant bidirectional effects between negative urgency scores and number of alcohol sips **(Table S12**). Negative urgency scores at year 2 significantly predicted alcohol sipping at year 4 (b = 0.27, p < 0.001), while alcohol sipping at year 2 predicted negative urgency scores at year 4 (b = 0.02, p < 0.001). There was a significant positive unidirectional effect of alcohol sipping at year 1 on negative urgency scores at year 2 (b = 0.02, p < 0.001), but the opposite relationship did not hold true. Moreover, bidirectional effects were observed between alcohol sipping and sensation seeking scores from year 2 to year 4 (b = 0.04, p < 0.05 for alcohol sipping at year 2 predicting sensation seeking at year 4; b = 0.12, p < 0.001 for the reverse). Additionally, BAS reward responsiveness and alcohol sipping affected each other simultaneously from year 2 to year 4 (b = 0.04, p < 0.05 for alcohol sipping at year 2 predicting BAS reward responsiveness at year 4; b = 0.05, p < 0.05 for the reverse). For all other outcomes, the bidirectional association was not observed in the data.

The RI-CLPM for negative urgency, sensation seeking, and BAS reward responsiveness achieved good fit to the data (CFI=0.946, TLI=0.912, RMSEA=0.017 for NU; CFI=0.969, TLI=0.950, RMSEA=0.015 for SS; CFI=0.944, TLI=0.909, RMSEA=0.017 for BAS RR).

Results show statistically significant bidirectional association between the number of alcohol sips and negative urgency over time (**Fig. 7**), that is the number of alcohol sips at year 2 positively impacting the negative urgency scores at year 4 (b = 0.08, p < 0.05) and negative urgency scores at year 2 showing a positive effect on the number of alcohol sips at year 4 (b = 0.09, p < 0.01). Additionally, negative urgency scores at baseline positively predicted negative urgency scores at year 2 (b = 0.14, p < 0.001), which further predicted negative urgency scores at year 4 (b = 0.25, p < 0.001). The number of alcohol sips at year 2 positively predicted the number of alcohol sips at year 4 (b = 0.45, p < 0.005). The number of alcohol sips and negative urgency scores covaried at year 4 (cov = 0.75, p < 0.05), suggesting that at year 4 an increased number of alcohol sips would be associated with greater negative urgency. There was evidence of stable, trait-like differences between persons for negative urgency scores (Var = 0.89, p < 0.001) and for the number of alcohol sips (Var = 1.42, p < 0.001). The models for sensation seeking and the BAS reward responsiveness did not show any bidirectional effects at any time point.

**Fig 7.**
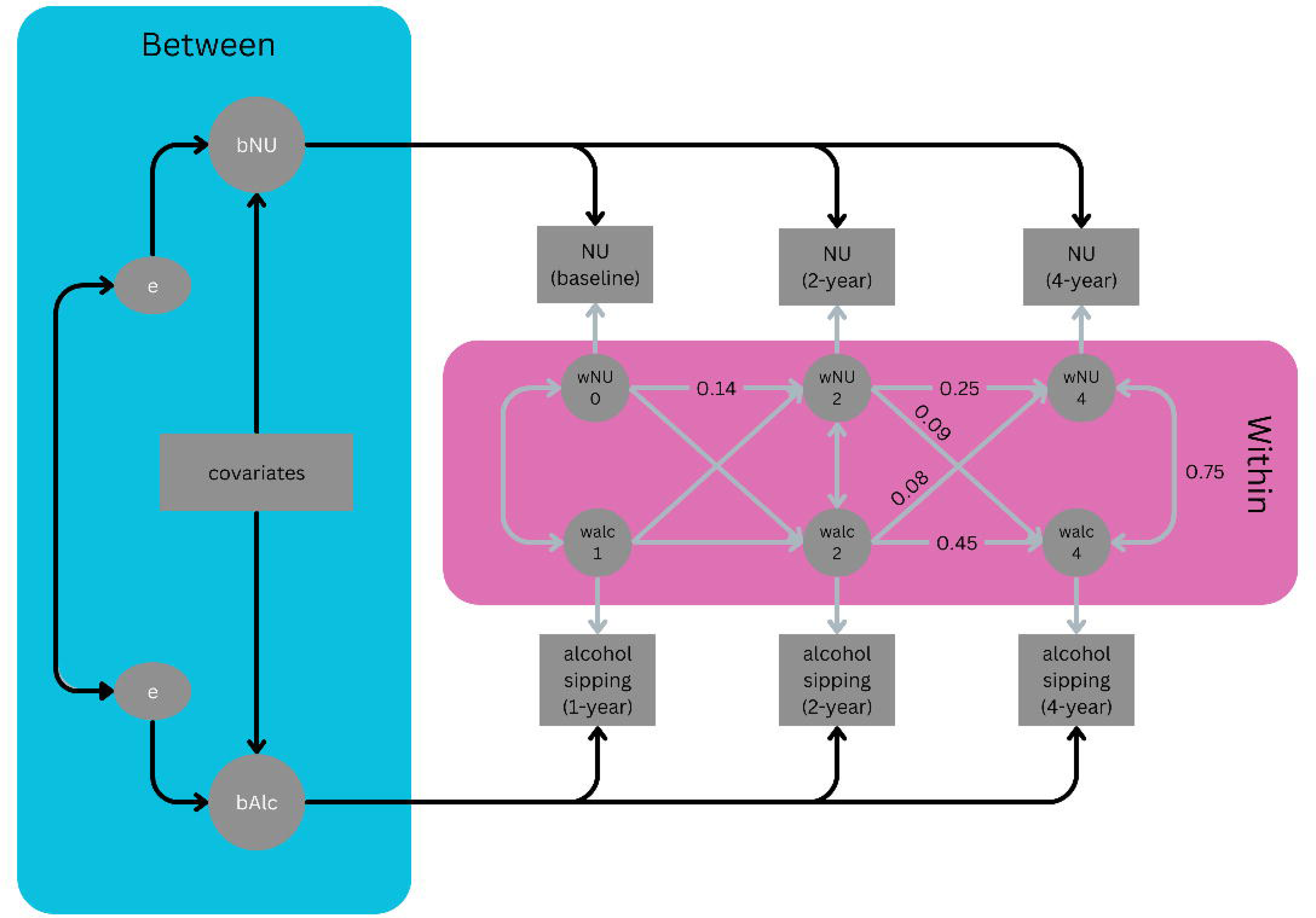
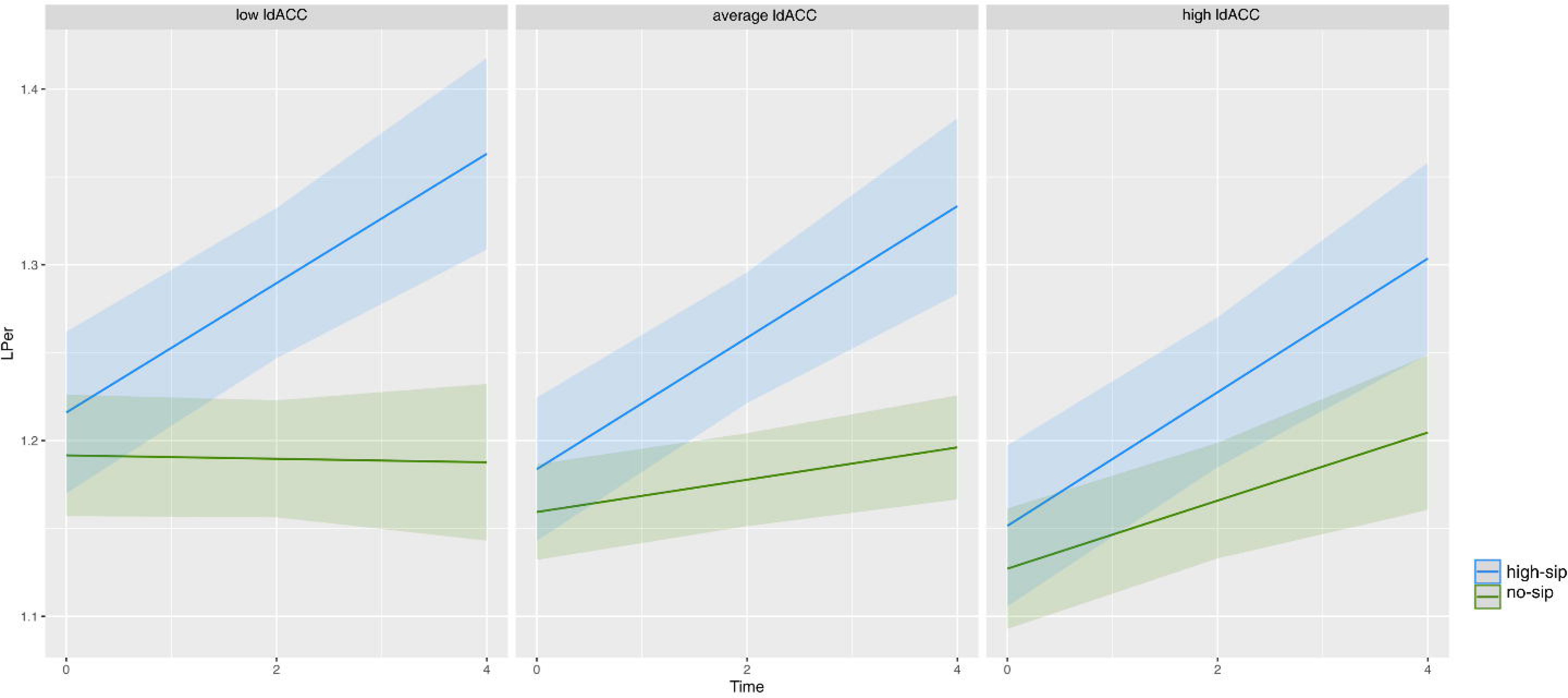
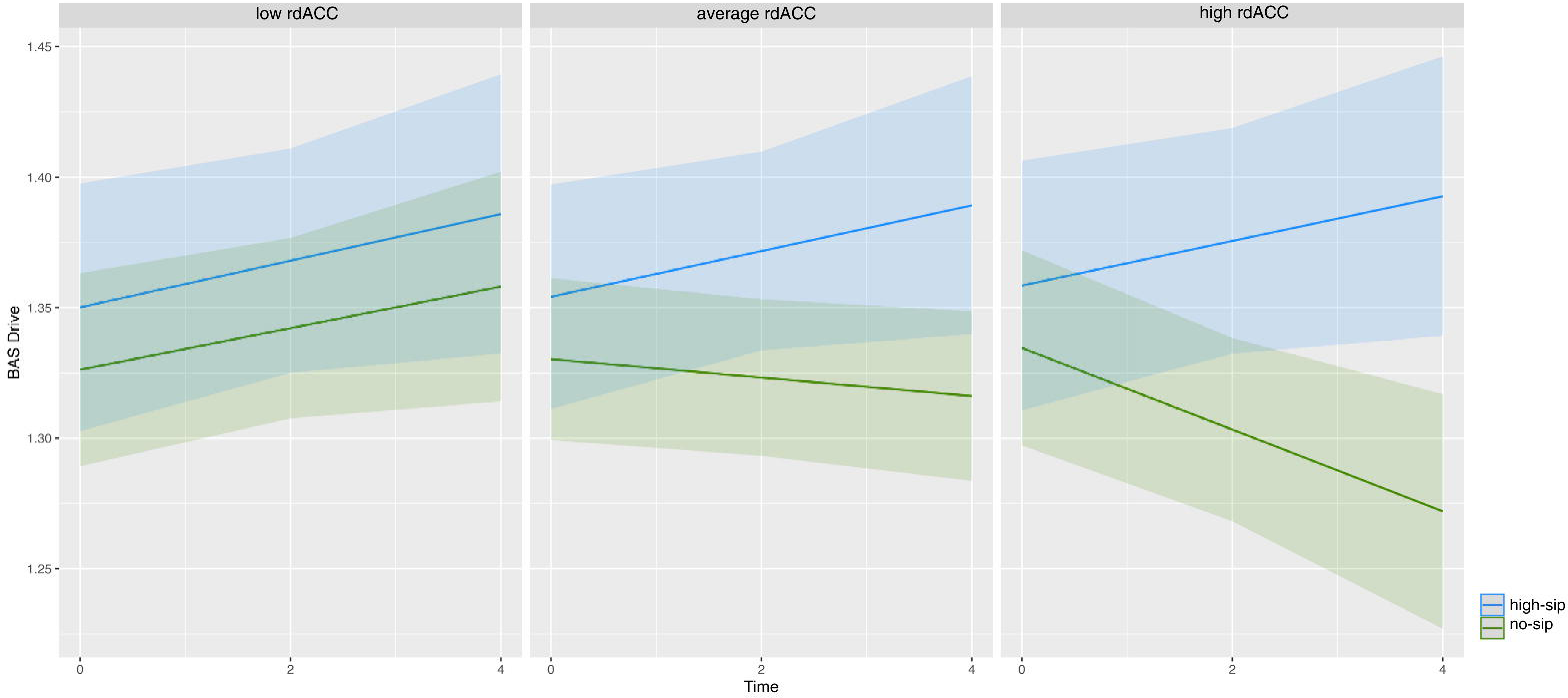
Graphic representation of the RI-CLPM showing bidirectional effects between negative urgency and alcohol sipping. *NU* denotes the observed negative urgency scores and *alc* denotes the observed number of alcohol sips. e is the error term of the random intercept.

## 4. Discussion

Our study conducted a comprehensive analysis of alcohol sipping behavior in children as they transition to adolescence. To our knowledge, this study is the first to examine the latent trajectories of alcohol sipping in a large cohort of young adolescents and, moreover, to extend the impact of these latent trajectories on psychopathology and personality traits. One of the novel findings in this investigation underscored the significant role of bilateral dACC baseline activation during the SST, indicative of response inhibition and cognitive control, in moderating the association between early alcohol sipping patterns and outcomes related to mental health or personality traits. Notably, reduced activation in the left dACC and increased activation in the right dACC during this task exhibited contrasting outcomes between the high-sip group and the no-sip group or low-sip group in the trajectory of outcome scores. Additionally, our exploration extended to assessing bidirectional influences between alcohol sipping and personality traits.

While only negative urgency scores consistently demonstrated bidirectional effects regardless of the statistical methodologies used, a more detailed exploration of these phenomena is pending further investigations, contingent upon the availability of new data waves in the future.

The ABCD cohort started reporting alcohol sipping at the age of 9-10 years old and our study examined data up to the children’s age of 13-14 years. It was not surprising to find that the percentage of adolescents reporting more alcohol sipping increased over time, while the percentage of adolescents not trying alcoholic beverages decreased over time. Alcohol exposure had an increasing trajectory in the transition from childhood to adulthood (Maggs & Schulenberg, 2004). The latent class mixed models determined three significant latent classes representing the alcohol sipping behavior of the ABCD cohort: a no-sip group, a high-sip group (increasing sipping over time) and a low-sip group (decreasing sipping behavior over time). Several longitudinal investigations have identified diverse patterns of alcohol consumption as adolescents progress into adulthood, including non/low use trajectories, developmentally limited trajectories, heavy-use trajectories, and late-onset trajectories (Su et al., 2019). The absence of a heavy-use trajectory in our cohort may be attributed to the age range of 9-14 years old for the study population. While a group with a late-onset trajectory was observed, the sample size was insufficient for inclusion in the present analysis.

There were significant differences between the personality trait scores between the high- sip group and the no-sip or low-sip groups. Individuals sipping less on average or not at all showed reduced scores for impulsivity, behavioral inhibition and behavioral activation over time. Impulsivity scores in adolescence predicted the later rates of alcohol use (Thomsen et al., 2018) and they contributed as a risk factor in the development of problematic substance use (Kaiser et al., 2016). Individuals with high positive urgency scores tended to focus on the positive aspects of drinking and downplay the negative consequences, sustaining elevated drinking levels over time (Cyders et al., 2010). In general, lack of planning might be associated with alcohol misuse and dependence (MacKillop et al., 2007), while lack of perseverance and BAS fun might be related to the amount of alcohol consumed (Coskunpinar, et al., 2013; Franken & Muris, 2006). The low-sip and no-sip groups showed decreased trajectories of all personality trait scores over time compared to the high-sip group, except the contrast between no- sip and high- sip groups on lack of planning. These findings collectively highlighted the multifaceted impact of personality traits on sipping behavior as participants transition from childhood to adolescence and its potential implications for approaching risky drinking events.

Among the mental health outcomes examined in our analysis depression scores showed the most interesting results. Depressed individuals were at higher risk for developing alcohol dependence (Boschloo et al., 2011) and on the other hand, AUD had a negative effect on the development of depressive disorder (Bruce et al., 2005; Hasin et al., 1996). While these extreme cases have not been observed in the ABCD cohort, it is possible that this is due to the predominantly healthy nature of the participants, with most not engaging in heavy drinking. The high-sip group showed a non-linear increase in depression scores over time, while the depression scores for the no-sip and low-sip groups initially increased over time, peaking around 2-year follow-up and then decreasing.

Reduced left dACC activation in the response inhibition tasks implied increased neural activity in the left dACC during unsuccessful inhibitory response, which was also apparent in error detection (Polli et al., 2005). Increased right dACC activation during the SST was related to increased right dACC neural activation during successful inhibitory response (correct stop) or to decreased right dACC activity during unsuccessful inhibitory control (incorrect stop) and during successful execution of a planned response (correct go). Overall, there was evidence showing that the trajectories of personality trait scores diverged significantly more over time for participants with lower left dACC and higher right dACC activation at baseline, with the non- sippers or low sippers presenting a decreasing trend over time while the high sippers showing an increasing trend. These findings suggested distinct bilateral patterns of activation in the dACC which may be indicative of differential cognitive processing or inhibitory control mechanisms between individuals with varying levels of alcohol consumption.

Previous studies showed that adolescents suffering from depression exhibited a notable inflexibility in the local efficiency of the right dACC (Ho et al., 2017, Lichtenstein et al., 2016). Furthermore, the reduced local efficiency of the dACC during a modified response inhibition task was significantly correlated with an earlier onset of depression. These findings align with previous research indicating that Major Depressive Disorder (MDD) may have an impact on the developmental trajectory of functional networks (Ho et al., 2017). In line with these findings, our results emphasized that depression scores were significantly moderated by bilateral dACC activation and alcohol sipping patterns. Participants in the no-sip group with higher left dACC and lower right dACC activation at baseline showed the greatest difference in depression levels when compared to the high-sip group across all SST contrasts. Additionally, the low-sip group showed lower depression levels relative to the high-sip group when participants showed higher left dACC or lower right dACC activation at baseline.

Our analysis showed significant bidirectional effects between alcohol sipping and negative urgency, sensation seeking and reward responsiveness from year 2 to year 4. Previous studies found that individuals with higher negative urgency scores were more likely to cope with negative affects by engaging in using alcohol (Hallihan et al., 2023). Negative urgency scores were associated with impaired control as well as coma-induced drinking behavior and other risky behaviors (McCarthy et al., 2017). Other studies showed that there was a bidirectional effect between negative urgency, sensation seeking and alcohol intake over time (Gmel et al., 2009; Kaiser et al., 2016; Stamates, 2019), which was also observed in our study involving the transition from childhood to adolescence. Sensation seeking, identified as a consistent trait in childhood, appeared to drive children under the age of 14 toward seeking out novel and exciting behaviors, including alcohol use (Cappelli et al., 2020). Furthermore, we also observed significant bidirectional effects between BAS reward responsiveness and alcohol sipping. All reciprocal effects were positive, indicating that an increase in alcohol consumption corresponded to higher scores in negative urgency, sensation seeking, or BAS reward responsiveness and vice versa. However, only negative urgency showed significant bidirectional effects when using a more complex model, the RI-CLPM, highlighting the complexity of our data.

Our study has several limitations. First, personality traits tend to be inherited to some degree and could have significant effects on lifetime outcomes, including psychopathology (Sanchez-Roige et al., 2018). In our analysis we did not account for any genetic factors which might play an important role in the development of personality traits and mental health outcomes in adolescence. Second, within the ABCD study, caregivers provide annual reports on the child’s behavior, including CBCL and PGBI scores, until the children reach the age of 12. This aspect may introduce a potential source of bias, as prior research has emphasized differences between parent and youth assessments of symptomatology (Chen et al., 2017). Third, the nature of the study was designed to cover the diverse adolescent population in the United States and given the age range of 9-14 years old, it is unsurprising that the majority of healthy children refrained from alcohol experimentation or heavy use. Nevertheless, our findings indicate that even minimal alcohol experimentation in early adolescence may influence later alcohol use, potentially impacting subsequent developments in personality traits and mental health outcomes.

In the subsequent phases of our study, exploring new latent classes of alcohol sipping will be a compelling avenue, particularly with the availability of future time waves. Our latent class mixed models revealed a fourth potential group characterized by sustained abstinence initially, followed by a consistent increase over time. Although this class lacked sufficient participants for reliability in the current analysis, future data could provide a more robust foundation for examination. Additionally, extending the analysis to encompass various substance uses, beyond alcohol alone, and exploring the impact of co-use on the discussed personality traits and mental health outcomes would offer more valuable insights into the development of adolescent substance use. Moreover, a more in-depth examination of the bidirectionality between alcohol sipping and personality traits and psychopathology awaits future investigations, contingent upon the availability of additional data waves and more substance experimentation as participants transition from adolescence to young adulthood.

