## Supplemental Materials for "Alcohol Sipping Patterns, Personality, and Psychopathology in Children: Moderating Effects of Dorsal Anterior Cingulate Cortex (dACC)"

**FIGURE S1 HERE**

**Fig S1. The trajectory of lack of perseverance scores for the high-sip vs no-sip group for low, average and high left dACC activation in the correct-stop-vs-incorrect-stop SST contrast.**

**FIGURE S2 HERE**

**Fig. S2. The trajectory of BAS drive scores for the high-sip vs no-sip group for low, average and high right dACC activation in the correct-stop-vs-incorrect-stop SST contrast.**

**Table S1. Summary table for the latent class mixed models**

| **Number of classes examined** | **loglik** | **Number of parameters** | **BIC** | **AIC** | **% class 1** | **% class 2** | **% class 3** | **% class 4** |
| --- | --- | --- | --- | --- | --- | --- | --- | --- |
| 1 | -996.05 | 28 | 2253.94 | 2048.09 | 100.00 |  |  |  |
| 2 | 2500.65 | 31 | -4711.4 | -4939.3 | 90.34 | 9.66 |  |  |
| 3 | 2544.28 | 34 | -4770.61 | -5020.57 | 5.34 | 84.22 | 10.44 |  |
| 4 | 2101.33 | 37 | -3856.66 | -4128.67 | 11.20 | 7.90 | 78.69 | 2.21 |

**Table S2a. Summary table for the effects of early alcohol patterns on negative urgency moderated by the dACC in the correct-stop-vs-incorrect stop SST contrast**

| **Negative Urgency** | **Estimate** | **St. Error** | **t value** | **p-value** |
| --- | --- | --- | --- | --- |
| (Intercept) | 1.79 | 0.08 | 21.83 | 0.00 |
| time | 0.05 | 0.02 | 2.32 | 0.02 |
| lc1 | 0.03 | 0.04 | 0.79 | 0.43 |
| lc2 | -0.08 | 0.03 | -3.24 | 0.00 |
| left dACC | 0.02 | 0.07 | 0.30 | 0.76 |
| right dACC | 0.04 | 0.07 | 0.54 | 0.59 |
| Family history of alcoholism | 0.04 | 0.01 | 2.88 | 0.00 |
| Sex | 0.10 | 0.01 | 7.87 | 0.00 |
| Race | 0.06 | 0.01 | 4.11 | 0.00 |
| SES | 0.02 | 0.02 | 0.98 | 0.33 |
| Age | 0.00 | 0.00 | 2.20 | 0.03 |
| time:lc1 | -0.08 | 0.03 | -2.38 | 0.02 |
| time:lc2 | -0.11 | 0.02 | -5.19 | 0.00 |
| time:lc1:left dACC | -0.38 | 0.22 | -1.70 | 0.09 |
| time:lc2:left dACC | -0.07 | 0.06 | -1.11 | 0.27 |
| time:lc1:right dACC | 0.27 | 0.23 | 1.20 | 0.23 |
| time:lc2:right dACC | 0.02 | 0.07 | 0.33 | 0.74 |
| Notes: **lc1 and lc2 are dummy variables, representing high-sip (lc1=0, lc2=0), low-sip (lc1=1, lc2=0) and no-sip (lc1=0, lc2=1).** | | | | |

**Table S2b. Summary table for the effects of early alcohol patterns on negative urgency moderated by the dACC in the correct-stop-vs-correct-go SST contrast**

| **Negative Urgency** | **Estimate** | **St. Error** | **t value** | **p-value** |
| --- | --- | --- | --- | --- |
| (Intercept) | 1.79 | 0.08 | 21.75 | 0.00 |
| time | 0.05 | 0.02 | 2.33 | 0.02 |
| lc1 | 0.03 | 0.04 | 0.76 | 0.45 |
| lc2 | -0.08 | 0.03 | -3.21 | 0.00 |
| left dACC | 0.06 | 0.08 | 0.69 | 0.49 |
| right dACC | -0.06 | 0.09 | -0.70 | 0.49 |
| Family History of Alcoholism | 0.04 | 0.01 | 2.87 | 0.00 |
| Sex | 0.10 | 0.01 | 7.94 | 0.00 |
| Race | 0.06 | 0.01 | 4.08 | 0.00 |
| SES | 0.02 | 0.02 | 0.99 | 0.32 |
| Age | 0.00 | 0.00 | 2.24 | 0.03 |
| time:lc1 | -0.07 | 0.04 | -1.99 | 0.047 |
| time:lc2 | -0.11 | 0.02 | -4.97 | 0.00 |
| time:lc1:left dACC | 0.08 | 0.25 | 0.32 | 0.75 |
| time:lc2:left dACC | -0.06 | 0.08 | -0.72 | 0.47 |
| time:lc1:right dACC | -0.12 | 0.28 | -0.43 | 0.67 |
| time:lc2:right dACC | 0.03 | 0.08 | 0.31 | 0.75 |
| Notes: **lc1 and lc2 are dummy variables, representing high-sip (lc1=0, lc2=0), low-sip (lc1=1, lc2=0) and no-sip (lc1=0, lc2=1).** | | | | |

**Table S3a. Summary table for the effects of early alcohol patterns on positive urgency moderated by the dACC in the correct-stop-vs-incorrect stop SST contrast**

| **Positive Urgency** | **Estimate** | **St. Error** | **t value** | **p-value** |
| --- | --- | --- | --- | --- |
| (Intercept) | 1.18 | 0.11 | 11.18 | 0.00 |
| time | 0.05 | 0.02 | 1.91 | 0.06 |
| lc1 | 0.03 | 0.05 | 0.52 | 0.60 |
| lc2 | -0.10 | 0.03 | -3.17 | 0.00 |
| left dACC | -0.10 | 0.09 | -1.08 | 0.28 |
| right dACC | 0.11 | 0.09 | 1.25 | 0.21 |
| Family History of Alcoholism | 0.03 | 0.02 | 1.52 | 0.13 |
| Sex | 0.15 | 0.02 | 10.01 | 0.00 |
| Race | 0.15 | 0.02 | 8.49 | 0.00 |
| SES | 0.10 | 0.02 | 4.52 | 0.00 |
| Age | 0.00 | 0.00 | -0.93 | 0.35 |
| time:lc1 | -0.12 | 0.04 | -3.02 | 0.00 |
| time:lc2 | -0.10 | 0.03 | -4.00 | 0.00 |
| time:lc1:left dACC | -0.14 | 0.28 | -0.49 | 0.63 |
| time:lc2:left dACC | -0.07 | 0.08 | -0.93 | 0.35 |
| time:lc1:right dACC | 0.02 | 0.29 | 0.09 | 0.93 |
| time:lc2:right dACC | 0.04 | 0.08 | 0.52 | 0.60 |
| Notes: **lc1 and lc2 are dummy variables, representing high-sip (lc1=0, lc2=0), low-sip (lc1=1, lc2=0) and no-sip (lc1=0, lc2=1).** | | | | |

**Table S3b. Summary table for the effects of early alcohol patterns on positive urgency moderated by the dACC in the correct-stop-vs-correct-go SST contrast**

| **Positive Urgency** | **Estimate** | **St. Error** | **t value** | **p-value** |
| --- | --- | --- | --- | --- |
| (Intercept) | 1.18 | 0.11 | 11.20 | 0.00 |
| time | 0.05 | 0.02 | 1.92 | 0.06 |
| lc1 | 0.03 | 0.05 | 0.51 | 0.61 |
| lc2 | -0.10 | 0.03 | -3.13 | 0.00 |
| left dACC | 0.08 | 0.10 | 0.79 | 0.43 |
| right dACC | -0.12 | 0.11 | -1.10 | 0.27 |
| Family History of Alcoholism | 0.02 | 0.02 | 1.48 | 0.14 |
| Sex | 0.16 | 0.02 | 10.07 | 0.00 |
| Race | 0.15 | 0.02 | 8.44 | 0.00 |
| SES | 0.10 | 0.02 | 4.50 | 0.00 |
| Age | 0.00 | 0.00 | -0.92 | 0.36 |
| time:lc1 | -0.10 | 0.04 | -2.25 | 0.02 |
| time:lc2 | -0.10 | 0.03 | -3.82 | 0.00 |
| time:lc1:left dACC | -0.36 | 0.31 | -1.15 | 0.25 |
| time:lc2:left dACC | -0.12 | 0.10 | -1.27 | 0.20 |
| time:lc1:right dACC | 0.04 | 0.35 | 0.10 | 0.92 |
| time:lc2:right dACC | 0.08 | 0.10 | 0.75 | 0.45 |
| Notes: **lc1 and lc2 are dummy variables, representing high-sip (lc1=0, lc2=0), low-sip (lc1=1, lc2=0) and no-sip (lc1=0, lc2=1).** | | | | |

**Table S4a. Summary table for the effects of early alcohol patterns on lack of planning moderated by the dACC in the correct-stop-vs-incorrect stop SST contrast**

| **Lack of Planning** | **Estimate** | **St. Error** | **t value** | **p-value** |
| --- | --- | --- | --- | --- |
| (Intercept) | 0.69 | 0.21 | 3.26 | 0.00 |
| time | 0.12 | 0.06 | 2.07 | 0.04 |
| lc1 | 0.20 | 0.12 | 1.61 | 0.11 |
| lc2 | -0.32 | 0.08 | -4.21 | 0.00 |
| left dACC | 0.05 | 0.21 | 0.23 | 0.82 |
| right dACC | -0.02 | 0.21 | -0.09 | 0.93 |
| Family History of Alcoholism | 0.14 | 0.04 | 3.75 | 0.00 |
| Sex | 0.32 | 0.04 | 9.00 | 0.00 |
| Race | -0.16 | 0.04 | -4.34 | 0.00 |
| SES | -0.02 | 0.04 | -0.36 | 0.72 |
| Age | 0.00 | 0.00 | 1.44 | 0.15 |
| time:lc1 | -0.29 | 0.10 | -2.90 | 0.00 |
| time:lc2 | -0.02 | 0.06 | -0.25 | 0.80 |
| time:lc1:left dACC | -0.39 | 0.66 | -0.59 | 0.55 |
| time:lc2:left dACC | 0.28 | 0.19 | 1.44 | 0.15 |
| time:lc1:right dACC | 0.39 | 0.67 | 0.58 | 0.56 |
| time:lc2:right dACC | -0.21 | 0.20 | -1.09 | 0.27 |
| Notes: **lc1 and lc2 are dummy variables, representing high-sip (lc1=0, lc2=0), low-sip (lc1=1, lc2=0) and no-sip (lc1=0, lc2=1).** | | | | |

**Table S4b. Summary table for the effects of early alcohol patterns on lack of planning moderated by the dACC in the correct-stop-vs-correct-go SST contrast**

| **Lack of Planning** | **Estimate** | **St. Error** | **t value** | **p-value** |
| --- | --- | --- | --- | --- |
| (Intercept) | 0.70 | 0.21 | 3.30 | 0.00 |
| time | 0.12 | 0.06 | 2.08 | 0.04 |
| lc1 | 0.20 | 0.12 | 1.61 | 0.11 |
| lc2 | -0.32 | 0.08 | -4.20 | 0.00 |
| left dACC | 0.03 | 0.24 | 0.12 | 0.91 |
| right dACC | -0.11 | 0.25 | -0.42 | 0.68 |
| Family History of Alcoholism | 0.14 | 0.04 | 3.75 | 0.00 |
| Sex | 0.32 | 0.04 | 8.96 | 0.00 |
| Race | -0.16 | 0.04 | -4.38 | 0.00 |
| SES | -0.02 | 0.04 | -0.38 | 0.71 |
| Age | 0.00 | 0.00 | 1.47 | 0.14 |
| time:lc1 | -0.24 | 0.10 | -2.31 | 0.02 |
| time:lc2 | -0.02 | 0.06 | -0.30 | 0.77 |
| time:lc1:left dACC | 0.22 | 0.74 | 0.30 | 0.77 |
| time:lc2:left dACC | 0.30 | 0.23 | 1.32 | 0.19 |
| time:lc1:right dACC | -0.77 | 0.82 | -0.94 | 0.35 |
| time:lc2:right dACC | -0.28 | 0.24 | -1.17 | 0.24 |
| Notes: **lc1 and lc2 are dummy variables, representing high-sip (lc1=0, lc2=0), low-sip (lc1=1, lc2=0) and no-sip (lc1=0, lc2=1).** | | | | |

**Table S5a. Summary table for the effects of early alcohol patterns on lack of perseverance moderated by the dACC in the correct-stop-vs-incorrect stop SST contrast**

| **Lack of Perseverance** | **Estimate** | **St. Error** | **t value** | **p-value** |
| --- | --- | --- | --- | --- |
| (Intercept) | 0.95 | 0.06 | 14.67 | 0.00 |
| time | 0.08 | 0.01 | 5.51 | 0.00 |
| lc1 | 0.01 | 0.03 | 0.33 | 0.74 |
| lc2 | -0.02 | 0.02 | -1.26 | 0.21 |
| left dACC | -0.16 | 0.05 | -3.00 | 0.00 |
| right dACC | 0.12 | 0.05 | 2.16 | 0.03 |
| Family History of Alcoholism | 0.06 | 0.01 | 5.13 | 0.00 |
| Sex | 0.06 | 0.01 | 5.63 | 0.00 |
| Race | 0.05 | 0.01 | 4.29 | 0.00 |
| SES | 0.05 | 0.01 | 3.78 | 0.00 |
| Age | 0.00 | 0.00 | -0.09 | 0.93 |
| time:lc1 | -0.06 | 0.02 | -2.57 | 0.01 |
| time:lc2 | -0.06 | 0.02 | -3.64 | 0.00 |
| time:lc1:left dACC | 0.11 | 0.17 | 0.63 | 0.53 |
| time:lc2:left dACC | 0.10 | 0.05 | 2.01 | 0.04 |
| time:lc1:right dACC | -0.02 | 0.18 | -0.11 | 0.91 |
| time:lc2:right dACC | -0.08 | 0.05 | -1.57 | 0.12 |
| Notes: **lc1 and lc2 are dummy variables, representing high-sip (lc1=0, lc2=0), low-sip (lc1=1, lc2=0) and no-sip (lc1=0, lc2=1).** | | | | |

**Table S5b. Summary table for the effects of early alcohol patterns on lack of perseverance moderated by the dACC in the correct-stop-vs-correct-go SST contrast**

| **Lack of Perseverance** | **Estimate** | **St. Error** | **t value** | **p-value** |
| --- | --- | --- | --- | --- |
| (Intercept) | 0.96 | 0.06 | 14.79 | 0.00 |
| time | 0.08 | 0.01 | 5.52 | 0.00 |
| lc1 | 0.01 | 0.03 | 0.33 | 0.74 |
| lc2 | -0.02 | 0.02 | -1.28 | 0.20 |
| left dACC | -0.11 | 0.06 | -1.71 | 0.09 |
| right dACC | 0.04 | 0.07 | 0.57 | 0.57 |
| Family History of Alcoholism | 0.05 | 0.01 | 5.11 | 0.00 |
| Sex | 0.05 | 0.01 | 5.54 | 0.00 |
| Race | 0.05 | 0.01 | 4.33 | 0.00 |
| SES | 0.05 | 0.01 | 3.76 | 0.00 |
| Age | 0.00 | 0.00 | -0.11 | 0.91 |
| time:lc1 | -0.07 | 0.03 | -2.86 | 0.00 |
| time:lc2 | -0.06 | 0.02 | -3.70 | 0.00 |
| time:lc1:left dACC | -0.07 | 0.19 | -0.35 | 0.72 |
| time:lc2:left dACC | 0.10 | 0.06 | 1.68 | 0.09 |
| time:lc1:right dACC | 0.18 | 0.22 | 0.84 | 0.40 |
| time:lc2:right dACC | -0.09 | 0.06 | -1.49 | 0.14 |
| Notes: **lc1 and lc2 are dummy variables, representing high-sip (lc1=0, lc2=0), low-sip (lc1=1, lc2=0) and no-sip (lc1=0, lc2=1).** | | | | |

**Table S6a. Summary table for the effects of early alcohol patterns on sensation seeking moderated by the dACC in the correct-stop-vs-incorrect stop SST contrast**

| **Sensation Seeking** | **Estimate** | **St. Error** | **t value** | **p-value** |
| --- | --- | --- | --- | --- |
| (Intercept) | 7.67 | 0.33 | 23.09 | 0.00 |
| time | 0.16 | 0.06 | 2.62 | 0.01 |
| lc1 | -0.02 | 0.14 | -0.15 | 0.88 |
| lc2 | -0.77 | 0.09 | -8.63 | 0.00 |
| left dACC | -0.01 | 0.25 | -0.04 | 0.97 |
| right dACC | 0.12 | 0.25 | 0.47 | 0.64 |
| Family History of Alcoholism | 0.09 | 0.05 | 1.87 | 0.06 |
| Sex | 0.65 | 0.05 | 14.09 | 0.00 |
| Race | -0.23 | 0.05 | -4.29 | 0.00 |
| SES | -0.18 | 0.07 | -2.54 | 0.01 |
| Age | 0.02 | 0.00 | 5.39 | 0.00 |
| time:lc1 | -0.36 | 0.11 | -3.39 | 0.00 |
| time:lc2 | -0.16 | 0.07 | -2.47 | 0.01 |
| time:lc1:left dACC | -0.26 | 0.76 | -0.33 | 0.74 |
| time:lc2:left dACC | 0.44 | 0.21 | 2.10 | 0.04 |
| time:lc1:right dACC | -0.43 | 0.78 | -0.55 | 0.58 |
| time:lc2:right dACC | -0.49 | 0.21 | -2.28 | 0.02 |
| Notes: **lc1 and lc2 are dummy variables, representing high-sip (lc1=0, lc2=0), low-sip (lc1=1, lc2=0) and no-sip (lc1=0, lc2=1).** | | | | |

**Table S6b. Summary table for the effects of early alcohol patterns on sensation seeking moderated by the dACC in the correct-stop-vs-correct-go SST contrast**

| **Sensation Seeking** | **Estimate** | **St. Error** | **t value** | **p-value** |
| --- | --- | --- | --- | --- |
| (Intercept) | 7.66 | 0.33 | 23.03 | 0.00 |
| time | 0.16 | 0.06 | 2.62 | 0.01 |
| lc1 | -0.02 | 0.14 | -0.13 | 0.90 |
| lc2 | -0.77 | 0.09 | -8.62 | 0.00 |
| left dACC | -0.32 | 0.29 | -1.13 | 0.26 |
| right dACC | 0.41 | 0.30 | 1.35 | 0.18 |
| Family History of Alcoholism | 0.09 | 0.05 | 1.87 | 0.06 |
| Sex | 0.65 | 0.05 | 14.08 | 0.00 |
| Race | -0.23 | 0.05 | -4.29 | 0.00 |
| SES | -0.18 | 0.07 | -2.54 | 0.01 |
| Age | 0.02 | 0.00 | 5.42 | 0.00 |
| time:lc1 | -0.26 | 0.11 | -2.35 | 0.02 |
| time:lc2 | -0.17 | 0.07 | -2.46 | 0.01 |
| time:lc1:left dACC | -0.64 | 0.85 | -0.76 | 0.45 |
| time:lc2:left dACC | 0.33 | 0.25 | 1.34 | 0.18 |
| time:lc1:right dACC | -0.54 | 0.95 | -0.56 | 0.57 |
| time:lc2:right dACC | -0.34 | 0.26 | -1.31 | 0.19 |
| Notes: **lc1 and lc2 are dummy variables, representing high-sip (lc1=0, lc2=0), low-sip (lc1=1, lc2=0) and no-sip (lc1=0, lc2=1).** | | | | |

**Table S7a. Summary table for the effects of early alcohol patterns on BIS moderated by the dACC in the correct-stop-vs-incorrect stop SST contrast**

| **BIS** | **Estimate** | **St. Error** | **t value** | **p-value** |
| --- | --- | --- | --- | --- |
| (Intercept) | 2.45 | 0.08 | 30.39 | 0.00 |
| time | 0.03 | 0.02 | 1.78 | 0.08 |
| lc1 | 0.05 | 0.04 | 1.34 | 0.18 |
| lc2 | 0.01 | 0.02 | 0.58 | 0.56 |
| left dACC | -0.04 | 0.07 | -0.66 | 0.51 |
| right dACC | 0.10 | 0.07 | 1.52 | 0.13 |
| Family History of Alcoholism | 0.03 | 0.01 | 2.06 | 0.04 |
| Sex | -0.20 | 0.01 | -16.92 | 0.00 |
| Race | -0.05 | 0.01 | -3.42 | 0.00 |
| SES | -0.04 | 0.02 | -2.28 | 0.02 |
| Age | 0.00 | 0.00 | 3.21 | 0.00 |
| time:lc1 | -0.03 | 0.03 | -0.88 | 0.38 |
| time:lc2 | -0.10 | 0.02 | -4.98 | 0.00 |
| time:lc1:left dACC | 0.11 | 0.23 | 0.50 | 0.61 |
| time:lc2:left dACC | 0.04 | 0.06 | 0.57 | 0.57 |
| time:lc1:right dACC | -0.20 | 0.23 | -0.88 | 0.38 |
| time:lc2:right dACC | -0.09 | 0.06 | -1.42 | 0.16 |
| Notes: **lc1 and lc2 are dummy variables, representing high-sip (lc1=0, lc2=0), low-sip (lc1=1, lc2=0) and no-sip (lc1=0, lc2=1).** | | | | |

**Table S7b. Summary table for the effects of early alcohol patterns on BIS moderated by the dACC in the correct-stop-vs-correct-go SST contrast**

| **BIS** | **Estimate** | **St. Error** | **t value** | **p-value** |
| --- | --- | --- | --- | --- |
| (Intercept) | 2.45 | 0.08 | 30.28 | 0.00 |
| time | 0.03 | 0.02 | 1.79 | 0.07 |
| lc1 | 0.05 | 0.04 | 1.35 | 0.18 |
| lc2 | 0.01 | 0.02 | 0.60 | 0.55 |
| left dACC | 0.01 | 0.08 | 0.16 | 0.88 |
| right dACC | 0.01 | 0.08 | 0.17 | 0.86 |
| Family History of Alcoholism | 0.03 | 0.01 | 2.01 | 0.04 |
| Sex | -0.20 | 0.01 | -16.83 | 0.00 |
| Race | -0.05 | 0.01 | -3.41 | 0.00 |
| SES | -0.04 | 0.02 | -2.25 | 0.02 |
| Age | 0.00 | 0.00 | 3.22 | 0.00 |
| time:lc1 | -0.03 | 0.03 | -0.91 | 0.36 |
| time:lc2 | -0.10 | 0.02 | -4.93 | 0.00 |
| time:lc1:left dACC | -0.09 | 0.25 | -0.35 | 0.73 |
| time:lc2:left dACC | -0.05 | 0.07 | -0.63 | 0.53 |
| time:lc1:right dACC | 0.10 | 0.28 | 0.35 | 0.73 |
| time:lc2:right dACC | 0.05 | 0.08 | 0.59 | 0.55 |
| Notes: **lc1 and lc2 are dummy variables, representing high-sip (lc1=0, lc2=0), low-sip (lc1=1, lc2=0) and no-sip (lc1=0, lc2=1).** | | | | |

**Table S8a. Summary table for the effects of early alcohol patterns on BAS reward responsiveness moderated by the dACC in the correct-stop-vs-incorrect stop SST contrast**

| **BAS reward responsiveness** | **Estimate** | **St. Error** | **t value** | **p-value** |
| --- | --- | --- | --- | --- |
| (Intercept) | 8.40 | 0.28 | 30.23 | 0.00 |
| time | -0.52 | 0.06 | -8.20 | 0.00 |
| lc1 | 0.17 | 0.13 | 1.31 | 0.19 |
| lc2 | 0.01 | 0.08 | 0.09 | 0.93 |
| left dACC | 0.22 | 0.22 | 1.00 | 0.32 |
| right dACC | -0.10 | 0.23 | -0.43 | 0.67 |
| Family History of Alcoholism | -0.02 | 0.04 | -0.56 | 0.57 |
| Sex | 0.15 | 0.04 | 3.78 | 0.00 |
| Race | 0.23 | 0.05 | 5.08 | 0.00 |
| SES | 0.03 | 0.06 | 0.46 | 0.65 |
| Age | 0.00 | 0.00 | 0.46 | 0.64 |
| time:lc1 | -0.11 | 0.11 | -0.99 | 0.32 |
| time:lc2 | -0.17 | 0.07 | -2.45 | 0.01 |
| time:lc1:left dACC | 0.39 | 0.76 | 0.52 | 0.60 |
| time:lc2:left dACC | -0.26 | 0.21 | -1.22 | 0.22 |
| time:lc1:right dACC | -0.51 | 0.77 | -0.66 | 0.51 |
| time:lc2:right dACC | 0.02 | 0.21 | 0.12 | 0.91 |
| Notes: **lc1 and lc2 are dummy variables, representing high-sip (lc1=0, lc2=0), low-sip (lc1=1, lc2=0) and no-sip (lc1=0, lc2=1).** | | | | |

**Table S8b. Summary table for the effects of early alcohol patterns on BAS reward responsiveness moderated by the dACC in the correct-stop-vs-correct-go SST contrast**

| **BAS reward responsiveness** | **Estimate** | **St. Error** | **t value** | **p-value** |
| --- | --- | --- | --- | --- |
| (Intercept) | 8.42 | 0.28 | 30.24 | 0.00 |
| time | -0.52 | 0.06 | -8.19 | 0.00 |
| lc1 | 0.17 | 0.13 | 1.32 | 0.19 |
| lc2 | 0.01 | 0.08 | 0.11 | 0.91 |
| left dACC | 0.06 | 0.26 | 0.24 | 0.81 |
| right dACC | -0.22 | 0.27 | -0.80 | 0.42 |
| Family History of Alcoholism | -0.02 | 0.04 | -0.56 | 0.58 |
| Sex | 0.15 | 0.04 | 3.72 | 0.00 |
| Race | 0.23 | 0.05 | 5.05 | 0.00 |
| SES | 0.03 | 0.06 | 0.43 | 0.67 |
| Age | 0.00 | 0.00 | 0.45 | 0.65 |
| time:lc1 | -0.10 | 0.11 | -0.91 | 0.37 |
| time:lc2 | -0.17 | 0.07 | -2.46 | 0.01 |
| time:lc1:left dACC | 0.23 | 0.84 | 0.28 | 0.78 |
| time:lc2:left dACC | -0.18 | 0.25 | -0.72 | 0.47 |
| time:lc1:right dACC | -0.38 | 0.94 | -0.40 | 0.69 |
| time:lc2:right dACC | 0.26 | 0.26 | 0.97 | 0.33 |
| Notes: **lc1 and lc2 are dummy variables, representing high-sip (lc1=0, lc2=0), low-sip (lc1=1, lc2=0) and no-sip (lc1=0, lc2=1).** | | | | |

**Table S9a. Summary table for the effects of early alcohol patterns on BAS fun moderated by the dACC in the correct-stop-vs-incorrect stop SST contrast**

| **BAS fun** | **Estimate** | **St. Error** | **t value** | **p-value** |
| --- | --- | --- | --- | --- |
| (Intercept) | 2.06 | 0.08 | 26.69 | 0.00 |
| time | -0.07 | 0.02 | -4.01 | 0.00 |
| lc1 | 0.03 | 0.04 | 0.85 | 0.39 |
| lc2 | -0.07 | 0.02 | -3.15 | 0.00 |
| left dACC | -0.06 | 0.06 | -0.92 | 0.36 |
| right dACC | 0.03 | 0.06 | 0.43 | 0.67 |
| Family History of Alcoholism | 0.01 | 0.01 | 0.95 | 0.34 |
| Sex | 0.07 | 0.01 | 6.05 | 0.00 |
| Race | 0.04 | 0.01 | 2.90 | 0.00 |
| SES | 0.04 | 0.02 | 2.71 | 0.01 |
| Age | 0.00 | 0.00 | -0.19 | 0.85 |
| time:lc1 | -0.11 | 0.03 | -3.78 | 0.00 |
| time:lc2 | -0.11 | 0.02 | -5.94 | 0.00 |
| time:lc1:left dACC | 0.15 | 0.20 | 0.77 | 0.44 |
| time:lc2:left dACC | 0.17 | 0.06 | 3.10 | 0.00 |
| time:lc1:right dACC | -0.22 | 0.20 | -1.11 | 0.27 |
| time:lc2:right dACC | -0.15 | 0.06 | -2.61 | 0.01 |
| Notes: **lc1 and lc2 are dummy variables, representing high-sip (lc1=0, lc2=0), low-sip (lc1=1, lc2=0) and no-sip (lc1=0, lc2=1).** | | | | |

**Table S9b. Summary table for the effects of early alcohol patterns on BAS fun moderated by the dACC in the correct-stop-vs-correct-go SST contrast**

| **BAS fun** | **Estimate** | **St. Error** | **t value** | **p-value** |
| --- | --- | --- | --- | --- |
| (Intercept) | 2.07 | 0.08 | 26.75 | 0.00 |
| time | -0.07 | 0.02 | -3.99 | 0.00 |
| lc1 | 0.03 | 0.04 | 0.88 | 0.38 |
| lc2 | -0.07 | 0.02 | -3.16 | 0.00 |
| left dACC | -0.17 | 0.07 | -2.35 | 0.02 |
| right dACC | 0.09 | 0.07 | 1.27 | 0.20 |
| Family History of Alcoholism | 0.01 | 0.01 | 1.04 | 0.30 |
| Sex | 0.06 | 0.01 | 5.93 | 0.00 |
| Race | 0.04 | 0.01 | 2.87 | 0.00 |
| SES | 0.04 | 0.02 | 2.67 | 0.01 |
| Age | 0.00 | 0.00 | -0.20 | 0.84 |
| time:lc1 | -0.09 | 0.03 | -2.99 | 0.00 |
| time:lc2 | -0.11 | 0.02 | -6.05 | 0.00 |
| time:lc1:left dACC | 0.47 | 0.22 | 2.11 | 0.03 |
| time:lc2:left dACC | 0.16 | 0.07 | 2.40 | 0.02 |
| time:lc1:right dACC | -0.73 | 0.25 | -2.97 | 0.00 |
| time:lc2:right dACC | -0.13 | 0.07 | -1.88 | 0.06 |
| Notes: **lc1 and lc2 are dummy variables, representing high-sip (lc1=0, lc2=0), low-sip (lc1=1, lc2=0) and no-sip (lc1=0, lc2=1).** | | | | |

**Table S10a. Summary table for the effects of early alcohol patterns on BAS drive moderated by the dACC in the correct-stop-vs-incorrect stop SST contrast**

| **BAS drive** | **Estimate** | **St. Error** | **t value** | **p-value** |
| --- | --- | --- | --- | --- |
| (Intercept) | 1.09 | 0.06 | 16.83 | 0.00 |
| time | 0.02 | 0.01 | 1.32 | 0.19 |
| lc1 | 0.00 | 0.03 | 0.02 | 0.98 |
| lc2 | -0.02 | 0.02 | -1.26 | 0.21 |
| left dACC | 0.01 | 0.05 | 0.23 | 0.82 |
| right dACC | 0.02 | 0.05 | 0.40 | 0.69 |
| Family History of Alcoholism | -0.03 | 0.01 | -2.85 | 0.00 |
| Sex | 0.10 | 0.01 | 10.52 | 0.00 |
| Race | 0.09 | 0.01 | 8.20 | 0.00 |
| SES | 0.05 | 0.01 | 3.65 | 0.00 |
| Age | 0.00 | 0.00 | 5.37 | 0.00 |
| time:lc1 | -0.02 | 0.02 | -0.92 | 0.36 |
| time:lc2 | -0.02 | 0.02 | -1.61 | 0.11 |
| time:lc1:left dACC | 0.05 | 0.16 | 0.28 | 0.78 |
| time:lc2:left dACC | 0.07 | 0.05 | 1.57 | 0.12 |
| time:lc1:right dACC | -0.04 | 0.16 | -0.22 | 0.83 |
| time:lc2:right dACC | -0.12 | 0.05 | -2.51 | 0.01 |
| Notes: **lc1 and lc2 are dummy variables, representing high-sip (lc1=0, lc2=0), low-sip (lc1=1, lc2=0) and no-sip (lc1=0, lc2=1).** | | | | |

**Table S10b. Summary table for the effects of early alcohol patterns on BAS drive moderated by the dACC in the correct-stop-vs-correct-go SST contrast**

| **BAS drive** | **Estimate** | **St. Error** | **t value** | **p-value** |
| --- | --- | --- | --- | --- |
| (Intercept) | **1.09** | 0.06 | 16.79 | 0.00 |
| time | 0.02 | 0.01 | 1.33 | 0.18 |
| lc1 | 0.00 | 0.03 | 0.04 | 0.97 |
| lc2 | -0.02 | 0.02 | -1.23 | 0.22 |
| right dACC | 0.00 | 0.04 | 0.01 | 0.99 |
| Family History of Alcoholism | **-0.03** | 0.01 | -2.87 | 0.00 |
| Sex | **0.09** | 0.01 | 10.48 | 0.00 |
| Race | **0.08** | 0.01 | 8.19 | 0.00 |
| SES | **0.05** | 0.01 | 3.66 | 0.00 |
| Age | **0.00** | 0.00 | 5.39 | 0.00 |
| time:lc1 | -0.02 | 0.03 | -0.78 | 0.43 |
| time:lc2 | -0.02 | 0.02 | -1.37 | 0.17 |
| time:lc1:left dACC | 0.28 | 0.18 | 1.52 | 0.13 |
| time:lc2:left dACC | **0.12** | 0.06 | 2.11 | 0.04 |
| time:lc1:right dACC | **-0.32** | 0.20 | -1.56 | 0.12 |
| time:lc2:right dACC | **-0.15** | 0.06 | -2.64 | 0.01 |
| Notes: **lc1 and lc2 are dummy variables, representing high-sip (lc1=0, lc2=0), low-sip (lc1=1, lc2=0) and no-sip (lc1=0, lc2=1).** | | | | |

**Table S11a. Summary table for the effects of early alcohol patterns on depression moderated by the dACC in the correct-stop-vs-incorrect stop SST contrast**

| **Depression** | **Estimate** | **St. Error** | **t value** | **p-value** |
| --- | --- | --- | --- | --- |
| (Intercept) | **-0.73** | 0.23 | -3.12 | 0.00 |
| time | **0.16** | 0.04 | 3.72 | 0.00 |
| lc1 | 0.14 | 0.10 | 1.30 | 0.19 |
| lc2 | -0.05 | 0.07 | -0.81 | 0.42 |
| right dACC | 0.00 | 0.18 | 0.01 | 0.99 |
| I(time^2) | -0.02 | 0.01 | -1.67 | 0.17 |
| left dACC | -0.06 | 0.18 | -0.33 | 0.74 |
| Family History of Alcoholism | **0.59** | 0.04 | 15.28 | 0.00 |
| Sex | 0.05 | 0.04 | 1.55 | 0.12 |
| Race | 0.01 | 0.04 | 0.34 | 0.73 |
| SES | **0.13** | 0.05 | 2.60 | 0.01 |
| Age | 0.00 | 0.00 | 1.08 | 0.28 |
| time:lc1 | 0.06 | 0.08 | 0.80 | 0.42 |
| time:lc2 | -0.00 | 0.05 | -0.05 | 0.96 |
| lc1:I(time^2) | **-0.04** | 0.02 | -1.85 | 0.06 |
| lc2:I(time^2) | **-0.02** | 0.01 | -1.84 | 0.07 |
| time:lc1:right dACC | **1.03** | 0.58 | 1.78 | 0.08 |
| time:lc2:right dACC | -0.23 | 0.15 | -1.54 | 0.13 |
| lc1:right dACC:I(time^2) | -0.14 | 0.16 | -0.92 | 0.36 |
| lc2:right dACC:I(time^2) | **0.08** | 0.04 | 2.09 | 0.04 |
| time:lc1:left dACC | -0.55 | 0.57 | -0.97 | 0.33 |
| time:lc2:left dACC | **0.35** | 0.15 | 2.40 | 0.02 |
| lc1:I(time^2):left dACC | -0.04 | 0.16 | -0.26 | 0.80 |
| lc2:I(time^2):left dACC | **-0.13** | 0.04 | -3.37 | 0.00 |
| Notes: **lc1 and lc2 are dummy variables, representing high-sip (lc1=0, lc2=0), low-sip (lc1=1, lc2=0) and no-sip (lc1=0, lc2=1).** | | | | |

**Table S11b. Summary table for the effects of early alcohol patterns on depression moderated by the dACC in the correct-stop-vs-correct-go SST contrast**

| **Depression** | **Estimate** | **St. Error** | **t value** | **p-value** |
| --- | --- | --- | --- | --- |
| (Intercept) | **-0.68** | 0.24 | -2.80 | 0.00 |
| time | **0.08** | 0.04 | 1.93 | 0.05 |
| lc1 | 0.13 | 0.10 | 1.34 | 0.18 |
| lc2 | -0.07 | 0.06 | -1.09 | 0.28 |
| right dACC | -0.39 | 0.21 | -1.90 | 0.06 |
| I(time^2) | -0.01 | 0.01 | -1.08 | 0.28 |
| left dACC | 0.16 | 0.20 | 0.82 | 0.41 |
| Family History of Alcoholism | **0.58** | 0.04 | 15.06 | 0.00 |
| Sex | **0.09** | 0.04 | 2.61 | 0.01 |
| Race | 0.01 | 0.040 | 0.30 | 0.77 |
| SES | **0.14** | 0.05 | 2.75 | 0.01 |
| Age | **0.00** | 0.00 | 1.71 | 0.09 |
| time:lc1 | 0.02 | 0.08 | 0.21 | 0.83 |
| time:lc2 | -0.01 | 0.05 | -0.25 | 0.80 |
| lc1:I(time^2) | -0.02 | 0.02 | -0.84 | 0.40 |
| lc2:I(time^2) | **-0.02** | 0.01 | -1.66 | 0.10 |
| time:lc1:right dACC | **2.43** | 0.74 | 3.28 | 0.00 |
| time:lc2:right dACC | -0.22 | 0.18 | -1.19 | 0.24 |
| lc1:right dACC:I(time^2) | **-0.34** | 0.20 | -1.688 | 0.09 |
| lc2:right dACC:I(time^2) | **0.08** | 0.05 | 1.731 | 0.08 |
| time:lc1:left dACC | **-1.92** | 0.67 | -2.860 | 0.00 |
| time:lc2:left dACC | 0.26 | 0.17 | 1.542 | 0.12 |
| lc1:I(time^2):left dACC | 0.17 | 0.19 | 0.911 | 0.36 |
| lc2:I(time^2):left dACC | **-0.09** | 0.05 | -1.965 | 0.049 |
| Notes: **lc1 and lc2 are dummy variables, representing high-sip (lc1=0, lc2=0), low-sip (lc1=1, lc2=0) and no-sip (lc1=0, lc2=1).** | | | | |

**Table S12. Summary table of bidirectional effects between the number of alcohol sips and negative urgency (NU), sensation seeking (SS) and BAS reward responsiveness (BAS RR).**

| **Model** |  | **Estimate** | **St. Error** | **t value** | **p-value** |
| --- | --- | --- | --- | --- | --- |
| NU 0 -> Alc 2 | (Intercept) | -1.90 | 0.72 | -2.63 | 0.01 |
|  | alc1 | 0.22 | 0.02 | 13.64 | 0.00 |
|  | nu.0bsl | 0.08 | 0.07 | 1.10 | 0.27 |
|  | Age | 0.02 | 0.01 | 3.23 | 0.00 |
|  | Race | -0.88 | 0.20 | -4.50 | 0.00 |
|  | SES | -0.73 | 0.16 | -4.67 | 0.00 |
|  | Sex | 0.05 | 0.12 | 0.40 | 0.69 |
|  | Family History of Alcoholism | 0.18 | 0.16 | 1.11 | 0.27 |
| Alc 1 -> NU 2 | (Intercept) | 1.01 | 0.10 | 9.90 | 0.00 |
|  | alc1 | 0.02 | 0.01 | 3.40 | 0.00 |
|  | nu.0bsl | 0.25 | 0.01 | 26.27 | 0.00 |
|  | Age | 0.01 | 0.00 | 5.18 | 0.00 |
|  | Race | 0.05 | 0.02 | 3.22 | 0.00 |
|  | SES | 0.03 | 0.02 | 1.53 | 0.13 |
|  | Sex | 0.07 | 0.02 | 4.94 | 0.00 |
|  | Family History of Alcoholism | 0.03 | 0.02 | 1.97 | 0.49 |
| NU 2 -> Alc 4 | (Intercept) | 1.23 | 0.51 | 2.42 | 0.02 |
|  | alc1 | 0.16 | 0.02 | 8.60 | 0.00 |
|  | nu.2yr | 0.27 | 0.07 | 3.97 | 0.00 |
|  | Age | 0.04 | 0.01 | 6.27 | 0.00 |
|  | Race | -0.62 | 0.12 | -5.03 | 0.00 |
|  | SES | -0.86 | 0.11 | -8.02 | 0.00 |
|  | Sex | -0.43 | 0.11 | -4.11 | 0.00 |
|  | Family History of Alcoholism | 0.01 | 0.12 | 0.07 | 0.94 |
| Alc 2 -> NU 4 | (Intercept) | 1.57 | 0.14 | 11.50 | 0.00 |
|  | nu.0bsl | 0.16 | 0.01 | 11.47 | 0.00 |
|  | alc2 | 0.02 | 0.01 | 3.23 | 0.00 |
|  | Age | 0.00 | 0.00 | 1.78 | 0.07 |
|  | Race | 0.04 | 0.02 | 1.68 | 0.09 |
|  | SES | 0.01 | 0.03 | 0.30 | 0.77 |
|  | Sex | -0.06 | 0.02 | -2.58 | 0.01 |
|  | Family History of Alcoholism | 0.02 | 0.02 | 0.70 | 0.49 |
| SS 0 -> Alc 2 | (Intercept) | -0.83 | 0.64 | -1.29 | 0.20 |
|  | alc1 | 0.24 | 0.02 | 14.33 | 0.00 |
|  | ss.0bsl | -0.01 | 0.02 | -0.41 | 0.68 |
|  | Age | 0.01 | 0.01 | 1.80 | 0.07 |
|  | Race | -1.07 | 0.18 | -5.90 | 0.00 |
|  | SES | -0.81 | 0.14 | -5.80 | 0.00 |
|  | Sex | 0.10 | 0.12 | 0.82 | 0.41 |
|  | Family History of Alcoholism | -0.00 | 0.15 | -0.01 | 0.99 |
| Alc 1 -> SS 2 | (Intercept) | 3.78 | 0.34 | 11.21 | 0.00 |
|  | alc1 | 0.07 | 0.02 | 4.22 | 0.00 |
|  | ss.0bsl | 0.42 | 0.01 | 46.19 | 0.00 |
|  | Age | 0.02 | 0.00 | 5.36 | 0.00 |
|  | Race | -0.19 | 0.06 | -3.45 | 0.00 |
|  | SES | -0.24 | 0.07 | -3.31 | 0.00 |
|  | Sex | 0.36 | 0.05 | 7.51 | 0.00 |
|  | Family History of Alcoholism | 0.05 | 0.05 | 1.05 | 0.30 |
| SS 2 -> Alc 4 | (Intercept) | 0.64 | 0.50 | 1.27 | 0.21 |
|  | alc1 | 0.15 | 0.02 | 8.40 | 0.00 |
|  | ss.2yr | 0.12 | 0.02 | 6.94 | 0.00 |
|  | Age | 0.04 | 0.01 | 6.24 | 0.00 |
|  | Race | -0.59 | 0.12 | -4.87 | 0.00 |
|  | SES | -0.76 | 0.11 | -7.03 | 0.00 |
|  | Sex | -0.52 | 0.11 | -4.95 | 0.00 |
|  | Family History of Alcoholism | -0.03 | 0.12 | -0.27 | 0.79 |
| Alc 2 -> SS 4 | (Intercept) | 4.22 | 0.50 | 8.53 | 0.00 |
|  | ss.0bsl | 0.34 | 0.01 | 24.03 | 0.00 |
|  | alc2 | 0.04 | 0.02 | 1.98 | 0.48 |
|  | Age | 0.02 | 0.01 | 3.23 | 0.00 |
|  | Race | -0.48 | 0.09 | -5.67 | 0.00 |
|  | SES | -0.06 | 0.11 | -0.54 | 0.59 |
|  | Sex | 0.20 | 0.08 | 2.69 | 0.01 |
|  | Family History of Alcoholism | -0.10 | 0.08 | -1.26 | 0.21 |
| BAS RR 0 -> Alc 2 | (Intercept) | -0.34 | 0.72 | -0.47 | 0.64 |
|  | alc1 | 0.21 | 0.02 | 13.21 | 0.00 |
|  | bas_rr_sum.0bsl | 0.01 | 0.02 | 0.29 | 0.77 |
|  | Age | 0.03 | 0.01 | 4.41 | 0.00 |
|  | Race | -0.75 | 0.21 | -3.64 | 0.00 |
|  | SES | -0.98 | 0.16 | -6.28 | 0.00 |
|  | Sex | -0.41 | 0.13 | -3.30 | 0.00 |
|  | Family History of Alcoholism | -0.60 | 0.17 | -3.60 | 0.00 |
| Alc 1 -> BAS RR 2 | (Intercept) | 5.88 | 0.32 | 18.57 | 0.00 |
|  | alc1 | 0.03 | 0.02 | 1.91 | 0.06 |
|  | bas_rr_sum.0bsl | 0.29 | 0.01 | 28.43 | 0.00 |
|  | Age | -0.00 | 0.00 | -0.53 | 0.60 |
|  | Race | 0.06 | 0.05 | 1.19 | 0.24 |
|  | SES | -0.12 | 0.07 | -1.77 | 0.08 |
|  | Sex | -0.00 | 0.05 | -0.04 | 0.97 |
|  | Family History of Alcoholism | 0.01 | 0.0 | 0.17 | 0.86 |
| BAS RR 2 -> Alc 4 | (Intercept) | 0.41 | 0.543 | 0.75 | 0.45 |
|  | alc1 | 0.16 | 0.02 | 8.48 | 0.00 |
|  | bas_rr_sum.2yr | 0.05 | 0.02 | 2.44 | 0.02 |
|  | Age | 0.05 | 0.01 | 6.79 | 0.00 |
|  | Race | -0.62 | 0.13 | -4.91 | 0.00 |
|  | SES | -0.70 | 0.11 | -6.14 | 0.00 |
|  | Sex | -0.44 | 0.11 | -4.09 | 0.00 |
|  | Family History of Alcoholism | 0.08 | 0.12 | 0.68 | 0.50 |
| Alc 2 -> BAS RR 4 | (Intercept) | 7.17 | 0.47 | 15.31 | 0.00 |
|  | bas_rr_sum.0bsl | 0.19 | 0.02 | 11.85 | 0.00 |
|  | alc2 | 0.04 | 0.02 | 1.99 | 0.047 |
|  | Age | 0.01 | 0.00 | 2.90 | 0.00 |
|  | Race | -0.13 | 0.08 | -1.53 | 0.13 |
|  | SES | -0.26 | 0.10 | -2.63 | 0.01 |
|  | Sex | -0.24 | 0.07 | -3.26 | 0.00 |
|  | Family History of Alcoholism | -0.06 | 0.08 | -0.76 | 0.45 |
